# SIPA1L2 controls trafficking and signaling of TrkB-containing amphisomes at presynaptic terminals

**DOI:** 10.1101/556266

**Authors:** Maria Andres-Alonso, Mohamed Raafet Ammar, Ioana Butnaru, Guilherme M. Gomes, Gustavo Acuna Sanhueza, Rajeev Raman, PingAn Yuanxiang, Maximilian Borgmeyer, Jeffrey Lopez-Rojas, Syed Ahsan Raza, Nicola Brice, Torben J. Hausrat, Tamar Macharadze, Silvia Diaz-Gonzalez, Mark Carlton, Antonio Virgilio Failla, Oliver Stork, Michaela Schweizer, Eckart D. Gundelfinger, Matthias Kneussel, Christina Spilker, Anna Karpova, Michael R. Kreutz

## Abstract

Amphisomes are transient organelles that derive from fusion of autophagosomes with late endosomes. They rapidly transform into degradative autolysosomes, whereas non-degradative roles of the autophagic pathway have been barely described. Here we show that in neurons BDNF/TrkB receptor bearing Rab7 / Light chain 3 (LC3) - positive amphisomes signal at presynaptic boutons during retrograde trafficking to the soma. Local signaling and inward transport essentially require the Rap GTPase-activating (RapGAP) protein SIPA1L2, which directly binds to TrkB and Snapin to connect TrkB-containing amphisomes to dynein. Association with LC3 regulates the RapGAP activity of SIPA1L2 and thereby retrograde trafficking. Following induction of presynaptic plasticity amphisomes dissociate from dynein at boutons, and this enables local signaling and promotes transmitter release. Accordingly, *sipa1l2* knockout mice show impaired BDNF-dependent presynaptic plasticity. Collectively, the data suggest that TrkB-signaling endosomes are in fact amphisomes that during retrograde transport have local signaling capacity in the context of presynaptic plasticity.

---

TrkB signaling endosomes that carry axonal neurotrophic signals are generated at axon terminals and constitute an important long-range retrograde signaling mechanism conveying information of synaptic activity (1). Upon binding to BDNF, BDNF/TrkB have been shown to be internalized either via pinocytosis (2, 3) or in a clathrin-dependent manner (4) to specialized vesicular compartments and sorted to diverse pathways by yet unknown mechanisms. TrkB signaling is required for different aspects of neuronal function including the expression of presynaptic LTP at mossy fiber (MF) synapses (5). By binding to different adaptor proteins, TrkB activates three major molecular cascades: the Ras-MAPK pathway, the phosphatidyl inositol-3 (PI3)-kinase cascade and the phospholipase C-γ1 pathway (6). Rap-1 based signaling ensures sustained ERK activation since prenylated Rap1 is associated to TrkB containing endosomes. These are long-lived and persist during transport from nerve terminals to the cell soma (7, 8 9,10).

SIPA1L2 (also known as SPAR2) is a member of the SIPA1L family of neuronal RapGAPs (Fig. S1) (11). The protein is most abundant in granule cells of the dentate gyrus (DG) and cerebellum and shows RapGAP activity for the small GTPases Rap1 and 2 (12). Here, we report that SIPA1L2 binds directly to TrkB, and links the receptor tyrosine kinase to a Dynein motor for retrograde trafficking via a direct interaction with the adaptor protein Snapin. Interestingly, SIPA1L2 concurrently associates via LC3b to Rab7-positive amphisomes and binding of LC3b promotes RapGAP activity. The amphisome trafficks retrogradely along axons, it stops at presynaptic boutons and both motility and signaling are controlled by SIPA1L2 whose RapGAP activity reduces the velocity of amphisome transport. Presynaptic long-term potentiation (LTP) induces a Protein kinase A (PKA)-dependent dissociation of the SIPA1L2/Snapin complex from Dynein intermediate chain (DIC). This increases synaptic dwelling time of the amphisome and PKA phosphorylation of SIPA1L2 reduces RapGAP activity and enables local TrkB signaling at boutons, which in turn promotes neurotransmitter release. *sipa1l2* knockout (ko) mice show impaired BDNF-dependent mossy fiber (MF) LTP and spatial pattern separation, which requires MF plasticity. Application of a TAT-peptide encompassing the binding region for TrkB in SIPA1L2 induces a similar phenotype *in vivo* and *in vitro* in wild-type (*wt*) mice like those observed in *sipa1l2* ko mice. Collectively the data suggest that retrograde axonal transport of BDNF/TrkB occurs in neuronal amphisomes that are involved in plasticity-relevant local signaling at presynaptic boutons.

## Results

### *sipa1l2*^−/−^ mice exhibit impaired BDNF-dependent MF plasticity and deficits in spatial pattern separation

To study the neuronal function of SIPA1L2 we generated *sipa1l2* ko mice (Fig. S2a-d). No major morphological abnormalities were observed in the cerebellum (Fig. S2e) and the DG (Fig. S2h-m), motor learning as well as coordination (Fig. S2f,g). The number of adult-born granule cells and measures of general DG excitability and postsynaptic function were all normal in *sipa1l2*^−/−^ mice (Fig. S2n,o, S3a-f). However, a significant deficit was found when we assessed MF LTP, a form of plasticity presynaptically expressed at MF boutons of DG granule cells (Fig. 1a,b). Accordingly, we also observed a strong impairment in spatial pattern separation, a cognitive process associated with proper DG function and MF LTP (13,14,15,16,17), that is responsible for the disambiguation and independent storage of similar memories (Fig 1c-e). No changes were found in novel object location and recognition paradigms that are sensitive to interruption of hippocampal CA1 synaptic function (Fig. 1f). Spatial pattern separation as well as MF LTP have been shown to depend on BDNF/TrkB signalling in the DG (5, 14) and we indeed found a reduction in MF LTP when we chelated endogenous BDNF during perfusion of slices with TrkB-Fc bodies (Fig. 1g,h), resembling the decline in late phase LTP in *sipa1l2*^−/−^ mice.

**Figure 1.**
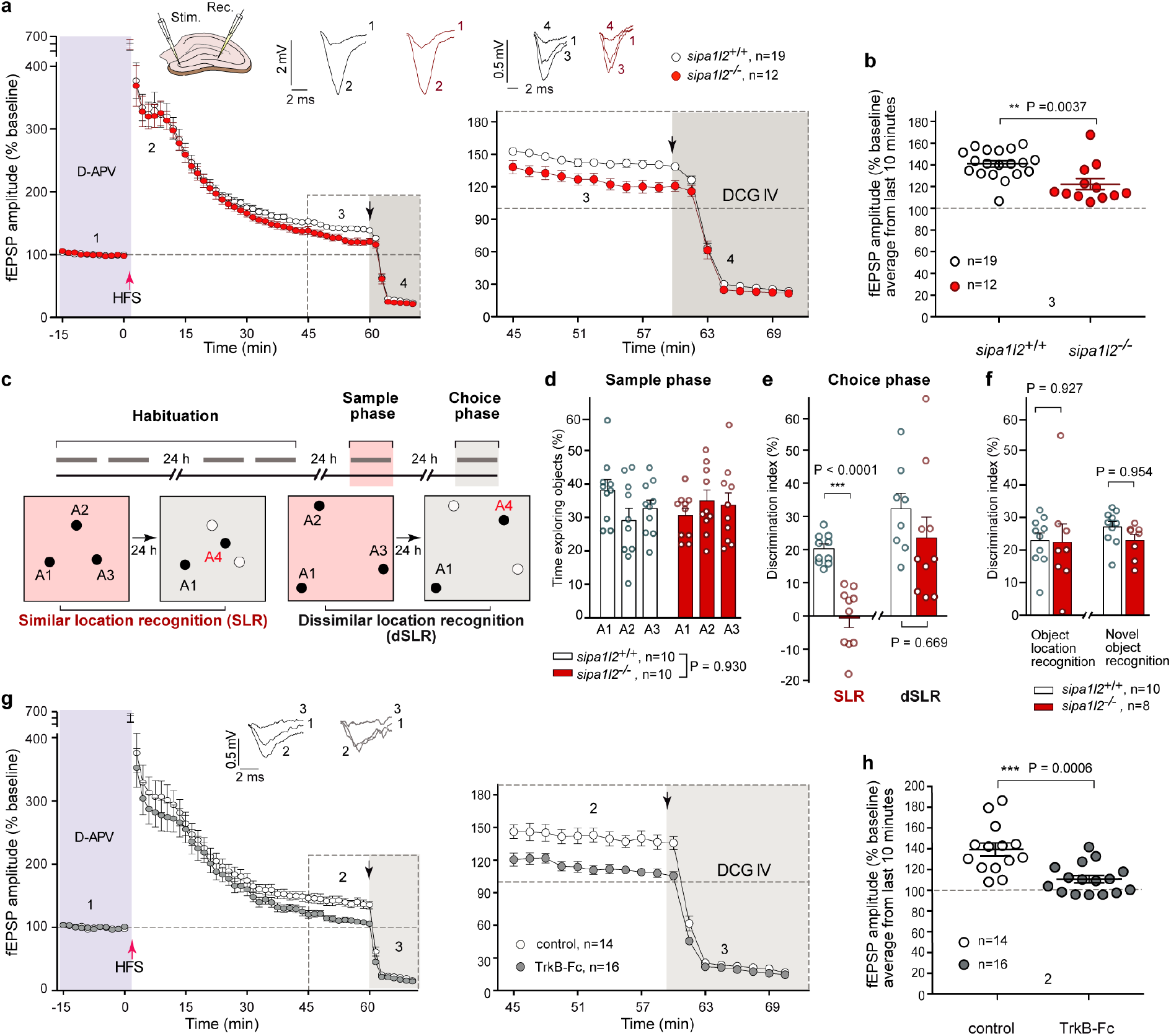
*sipa1l2*^−/−^ mice exhibit impaired MF plasticity and deficits in pattern separation. **a.** Presynaptic MF-LTP is reduced in *sipa1l2*^−/−^ acute slices. Left, average values of fEPSP amplitudes upon MF-LTP induction. Right, fEPSP amplitudes during the last 45-70 min following MF-LTP induction. **b**. Averaged fEPSP amplitudes recorded during the last 10 min of **a** (Mann-Whitney U test). **c.** Schematic representation of the spatial pattern separation behavior test. During the sample phase (red-shaded boxes) two groups of animals were trained in an arena: in the similar location recognition group (SLR), objects (A1-3) were placed closer to each other; and in the dissimilar location recognition group (dSLR), objects (A1-3) were placed further away from each other. During choice phase (grey-shaded boxes), a fourth object (A4) was placed between positions A2 and A3 in both groups. Animals from the SLR find the new object in A4 closer to the objects originally place in A2 and A3 and have a higher demand for pattern separation than those from the dSLR. Grey bars indicate 10 min interval. Filled circles (A1-4) represent object location. Open circles indicate absence of objects. **d.** Exploration time of *sipa1l2* wt and ko animals in A1-3 during the sample phase. Bars represent data as mean±S.E.M and open circles represent single subjects (two-wayANOVA). **e**. A significant reduction in the discrimination index during choice phase was found for *sipa1l2*^−/−^ mice in the SLR group but not in the dSLR (unpaired Student’s t test). **f**. Discrimination index of *sipa1l2* wt and ko animals during the novel object location recognition and object recognition test (unpaired Student’s t test). **g, h.** Left, average fEPSP amplitudes upon MF-LTP induction in control and BDNF-depleted slices (TrkB-Fc). Right, fEPSP amplitudes obtained during the last 45-70 min in left panel. **h.** Averaged fEPSP amplitudes obtained during the last 10 minutes of (**g**) (Mann-Whitney U test). Bars and error bars represent data as mean±S.E.M in all graphs. Circles represent mean values of individual subjects (d,e,f) or slices (b,h).

### The cytoplasmic domain of TrkB directly interacts with SIPA1L2

These experiments raised the question whether SIPA1L2 might be involved in presynaptic BDNF-TrkB signaling. BDNF/TrkB are internalized at distal axons and transported retrogradely in a Dynein-dependent manner as signaling endosomes (18, 19, 1) associated with Rap1 (6, 10). SIPA1L2 is prominently present at presynaptic terminals of primary neurons labeled by the presynaptic marker Symaptophysin 1 (Fig. 2a). We therefore wondered whether SIPA1L2 and TrkB might be part of a single protein complex *in vivo*. SIPA1L2 showed an overlapping distribution with Rap1 as well as with TrkB in subcellular fractionation experiments (Fig. S4a). Co-immunoprecipitation experiments from extracts of rat brain homogenates revealed that the protein is indeed present in immunoprecipitates generated with a TrkB-specific antibody and vice versa, TrkB could be precipitated with a SIPA1L2 antibody (Fig. 2b; Fig. S4b). Moreover, TrkB and SIPA1L2 are present in transport complexes immunoprecipitated by an antibody directed against DIC (Fig. 2c) and both proteins are localized to axon terminals in hippocampal primary neurons (Fig. 2d) where they were found in very close association as revealed by STED imaging (Fig. 2e,f).

**Figure 2.**
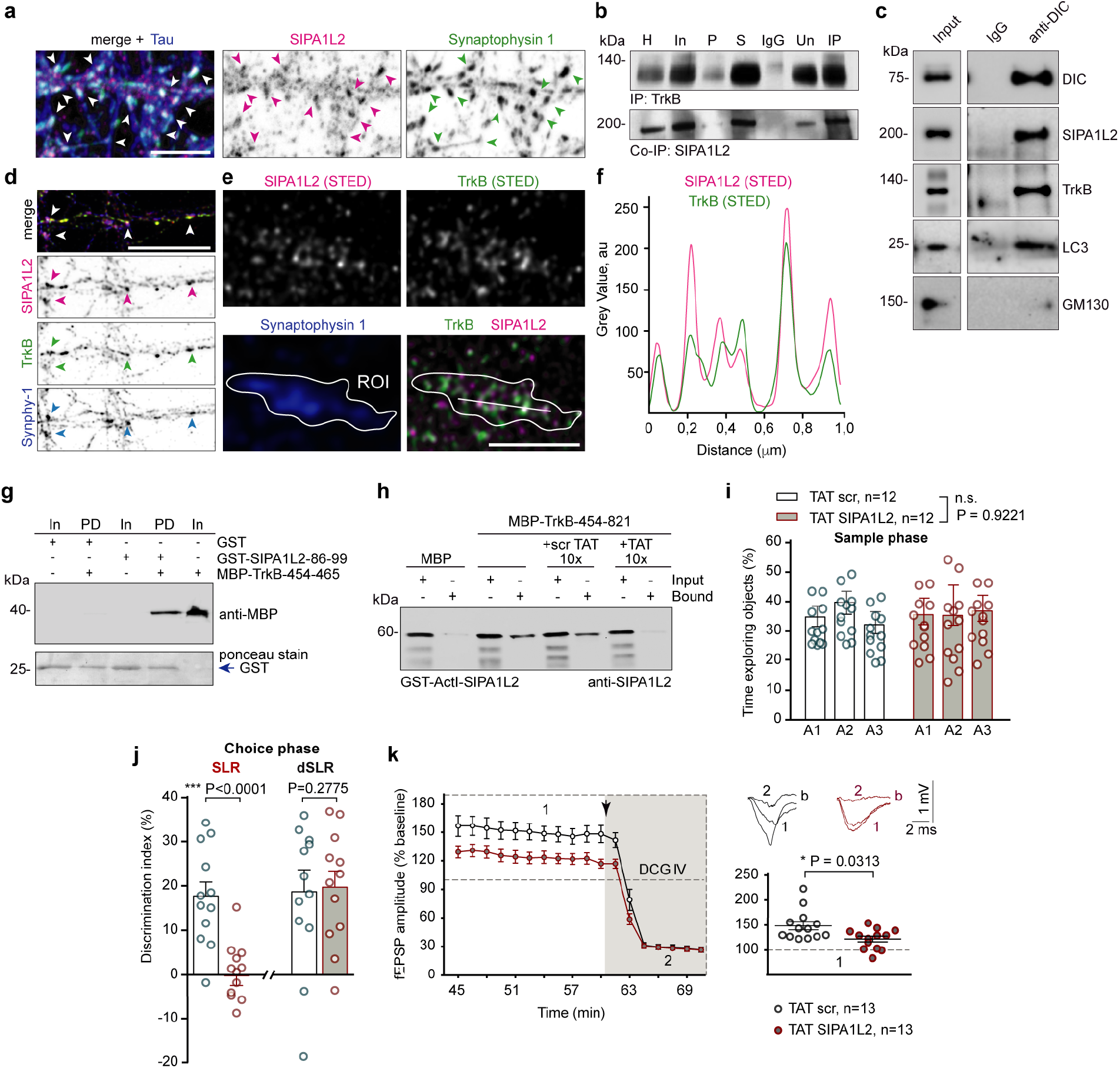
SIPA1L2 interaction with TrkB at presynaptic terminals is crucial for SIPA1L2 function in granule cells. **a.** Confocal images indicate the presence of SIPA1L2 at presynaptic boutons. Scale bar is 5 µm. **b**. Endogenous TrkB co-immunoprecipitates SIPA1L2 from cerebellar extracts (H-homogenate, In-input, P-pellet, S-supernatant, Un-unbound, IP-immunoprecipitation). **c.** Endogenous DIC co-immunoprecipitates SIPA1L2 together with TrkB and LC3 but not with GM130 from mouse forebrain. **d.** Confocal images show the co-localization of SIPA1L2 and TrkB at presynaptic boutons. Scale bar is 10 µm. **e, f.** STED images depicting the co-localization of SIPA1L2 and TrkB within a presynaptic bouton labeled by Synaptophysin-1. Line profiles (**f**) indicate relative intensities for STED channels along 1*μ*m. Scale bar is 1 *μ*m. **g**. Pull-down assay between the juxtamembrane region of TrkB and the 14aa of the binding interface in ActI-SIPA1L2 but not with GST. **h**. The interaction between TrkB and ActI-SIPA1L2 was prevented in a pull-down assay by addition of the 10X TAT binding peptide (TAT) but not 10X scrambled peptide (scrTAT). **i**. Time spent by *wt* mice injected with TAT-SIPA1L2 or TATscr peptides exploring objects (A1-3) during the sample phase (two-way ANOVA). **j**. Discrimination indexes obtained during the choice phase in SLR and dSLR groups injected with either TAT-SIPA1L2 or TATscr. Number of subjects is depicted in **i** (unpaired Student’s t test). **k**. fEPSP amplitudes recorded upon bath perfusion of the TAT-SIPA1L2 but not the TAT-scrambled peptide during the last 45 to 70 minutes after MF-LTP. Right, averaged values of fEPSP amplitudes during last 10 minutes of LTP recording (Mann-Whitney U test). Bars and error bars represent data as mean±S.E.M in all graphs. Circles represent mean values of individual subjects (**i,j**) or slices (**k**).

We next mapped the binding regions with Yeast-Two Hybrid (YTH) and found that the ActI-domain of SIPA1L2 interacts in the TrkB cytosolic domain with the first 12 juxtamembraneous amino acids (aa) (454-465) and the first 23 aa of the tyrosine kinase domain (537-559) (Fig. S4c-d). Bacterially expressed GST- and MBP-fusion proteins showed a direct interaction of cytoplasmic TrkB with the ActI but not the ActII domain of SIPA1L2 in pulldown assays (Fig. S4e). We next isolated 14 aa in SIPA1L2-ActI that are crucial for the association with TrkB (Fig. 2g; Fig. S4f, g, h). Of note, the binding region to TrkB is unique to SIPA1L2 when compared to other SIPA1L family members (Fig. S1b). Furthermore, a TAT-peptide containing this region competed for binding to TrkB in a pull-down assay (Fig. 2h). When we then infused this peptide into the DG of *wt* animals (Fig. S4i-j) we found diminished spatial pattern separation (Fig. 2i-j) similar to the one observed in *sipa1l2-/-* mice (Fig. 1e). In addition, bath perfusion of the TAT-peptide also reduced the amplitude of MF-LTP in acute slices (Fig. 2k). Thus, interruption of the SIPA1L2-TrkB interaction in *wt* mice mimicks the deficits of *sipa1l2*^−/−^ mice.

### SIPA1L2 interacts with Snapin and thereby enables TrkB retrograde trafficking

Dynein-dependent retrograde trafficking of TrkB-signaling endosomes in axons occurs in association with the adaptor protein Snapin, which recruits Dynein by interacting with DIC (19). However, whether Snapin interacts with its cargo directly or indirectly is still unclear. Heterologous co-immunoprecipitation experiments using tag-specific antibodies revealed that Snapin associates with the central region of SIPA1L2 encompassing the RapGAP and PDZ domains (Fig. 3a, Fig. S5a). The tagged RapGAP-PDZ domain co-recruits GFP-Snapin (Fig. 3b,c) and both proteins co-traffick in MRC5 cells (Fig. 3d). Snapin, TrkB and SIPA1L2 co-localize in axons of primary neurons (Fig. 3e) and live-imaging experiments using fluorescence-tagged proteins revealed a high percentage of mainly retrograde co-trajectories in axons (Fig. 3f-i, Fig. S5b). Of note, SIPA1L2 in dendrites remained immobile (Fig. S5c). Moreover, shRNA knockdown of Snapin reduced the motility of TrkB and SIPA1L2 and resulted in largely stationary complexes at the expense of less retrograde movement (Fig. 3j-k). Further, TrkB was found to remain largely stationary in *sipa1l2*^−/−^ neurons, supporting a role of SIPA1L2 in retrograde TrkB trafficking by mediating attachment to Dynein via Snapin (Fig. 3l-m).

**Figure 3.**
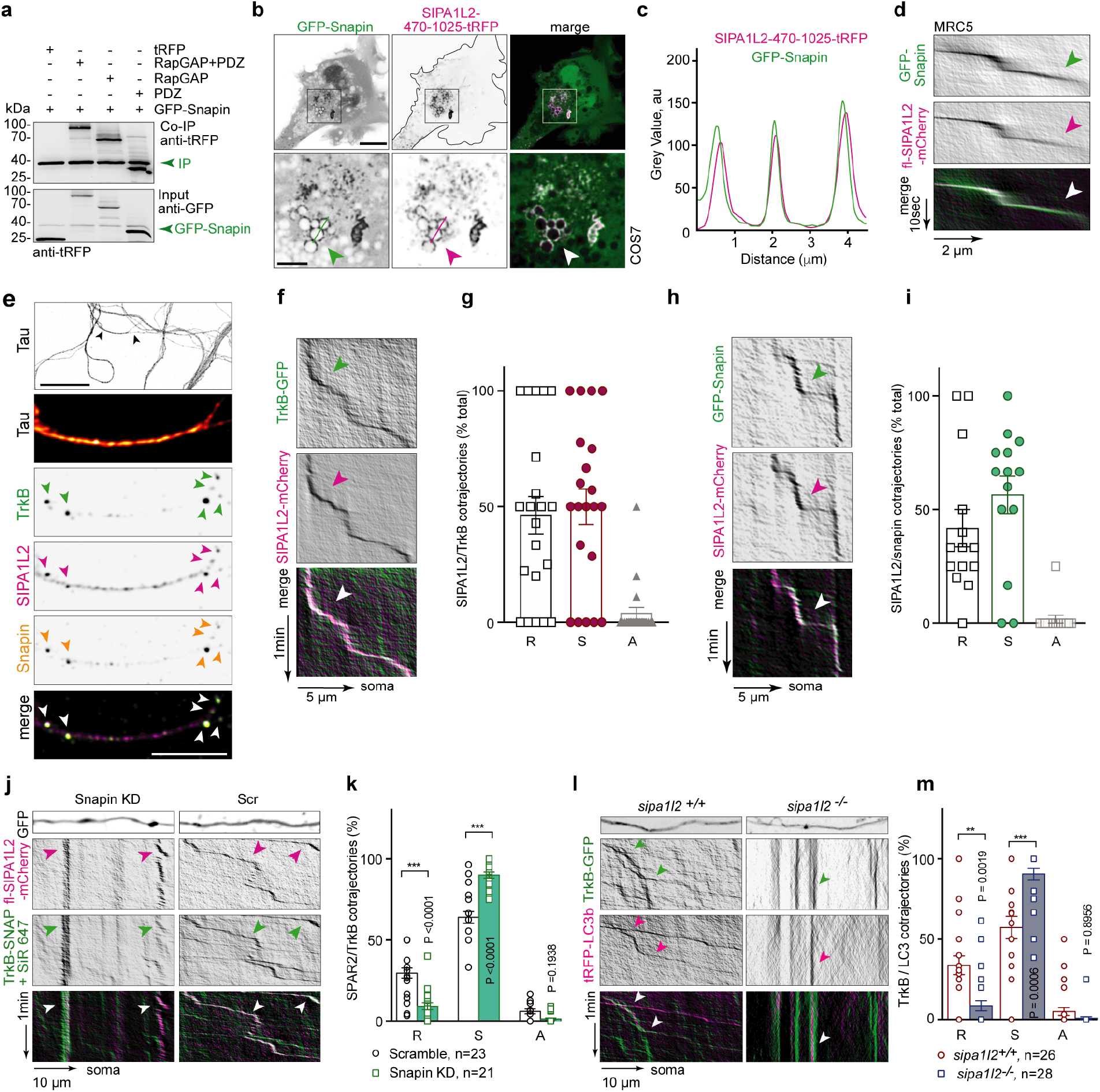
SIPA1L2 associates and co-trafficks with Snapin and TrkB and knock-down of Snapin immobilizes both, SIPA1L2 and TrkB. **a.** GFP-Snapin co-immunoprecipitates both RapGAP- (SIPA1L2-470-838-tRFP) and PDZ - SIPA1L2-tRFP from HEK293T cells extracts. **b,c**. Overexpression of SIPA1L2-470-1025-tRFP harboring RapGAP and PDZ domains in COS7 cells localizes to membranous structures and recruits GFP-Snapin as indicated with the line profile (**c**). Scale bars are 10 µm and 5 µm (inset). **d.** Representative kymographs obtained from MRC5 cells overexpressing WT-SIPA1L2-mCherry and GFP-Snapin. **e**. Confocal images from hippocampal neurons stained against Snapin, TrkB, SIPA1L2 and tau. Scale bars are 20 µm (upper panel) and 5 µm (lower panel). **f,g.** Representative kymographs prepared from axonal segments of primary neurons overexpressing TrkB-GFP and SIPA1L2-mCherry. In **g**, percentage of retrograde (R), stationary (S) and anterograde (A) axonal SIPA1L2/TrkB trajectories. **h,i.** Representative kymographs prepared from axonal segments of primary hippocampal neurons overexpressing GFP-Snapin and WT-SIPA1L2-mCherry. In **i**, percentage of cotrajectories as retrograde (R), stationary (S) and anterograde (A). **j,k.** Representative kymographs generated from axonal segments of neurons overexpressing either sh-based GFP-Snapin KD or control GFP-Scr together with fl-SIPA1L2-mCherry and TrkB-SNAP (+SiR647). Percentage of SIPA1L2/TrkB cotrajectories shows a significant reduction of the retrograde (R) trajectories with a correspondent enhancement of stationary (S) in neurons expressing GFP-Snapin KD as compared to the GFP-Scr (non-parametric Kolmogorov-Smirnov test; n is number of cells). **l,m**. Representative kymographs and quantification of percentage of TrkB-GFP/tRFP-LC3b cotrajectories in *sipa1l2*^−/−^ as compared to *sipa1l2*^+/+^ neurons (non-parametric Kolmogorov-Smirnov test). Bars and error bars represent data as mean±S.E.M. Symbols in columns represent values from single axons. Arrows indicate cotrajectories.

### Binding of LC3b to the RapGAP domain of SIPA1L2 modulates RapGAP activity

Autophagosomes recruit dynein and acquire Snapin by fusion with late endosomes and this fusion facilitates the retrograde trafficking of autophagosomes (20). In MRC5 cells we observed a prominent co-localization of SIPA1L2 with the autophagosomal marker LC3b but not with markers for lysosomes (LAMP1) and multivesicular bodies (CHMP4B) (Fig. S5d-e). Immunostaining of SIPA1L2, TrkB and LC3 in primary hippocampal neurons revealed extensive co-localization of all three proteins in axons (Fig. S5f) that was further increased in the presence of BDNF as quantified by Manders’ coefficient calculated for protein pairs (Fig. S5g,h). In addition, live imaging revealed largely retrograde cotrafficking of TrkB-GFP and tRFP-LC3b in a SIPA1L2-dependent manner (see above, Fig. 3l-m).

Interestinly, the RapGAP domain of SIPA1L2 contains a LIR-motif (FxxL-motif) for LC3-binding that is highly conserved in mammals and among all SIPA1L family members (21; Fig. S1b). In endogenous co-immunoprecipitation experiments performed an anti-DIC antibody to isolate transported complexes we found that LC3 co-precipitates with SIPA1L2 and TrkB (Fig. 2c). In addition, heterologous co-immunoprecipitation experiments conducted with tag-specific antibodies revealed an interaction of the RapGAP domain with LC3 in HEK293T cell lysates (Fig. 4a; Fig. S5a). Binding to LC3 was weaker when we expressed SIPA1L2 carrying point mutations within the LIR motif (Fig. 4b). Moreover, the RapGAP domain fused to GST efficiently pulled down endogenous LC3 from a rat brain lysate (Fig. 4c). Hence the catalytically active RapGAP domain of SIPA1L2 (23, aa 470-838; Fig. S1c, Fig. S5i,j) showed a direct interaction with bacterially expressed His-LC3b in a pull-down assay (Fig. 4d) and this interaction was not affected by inclusion of Snapin in the pulldown buffer (Fig. 4e). Reciprocally, the association with Snapin was not reduced by mutation of the LIR motif (Fig. 4f). Thus, it appears that both Snapin and LC3 can concomitantly bind to SIPA1L2.

**Figure 4.**
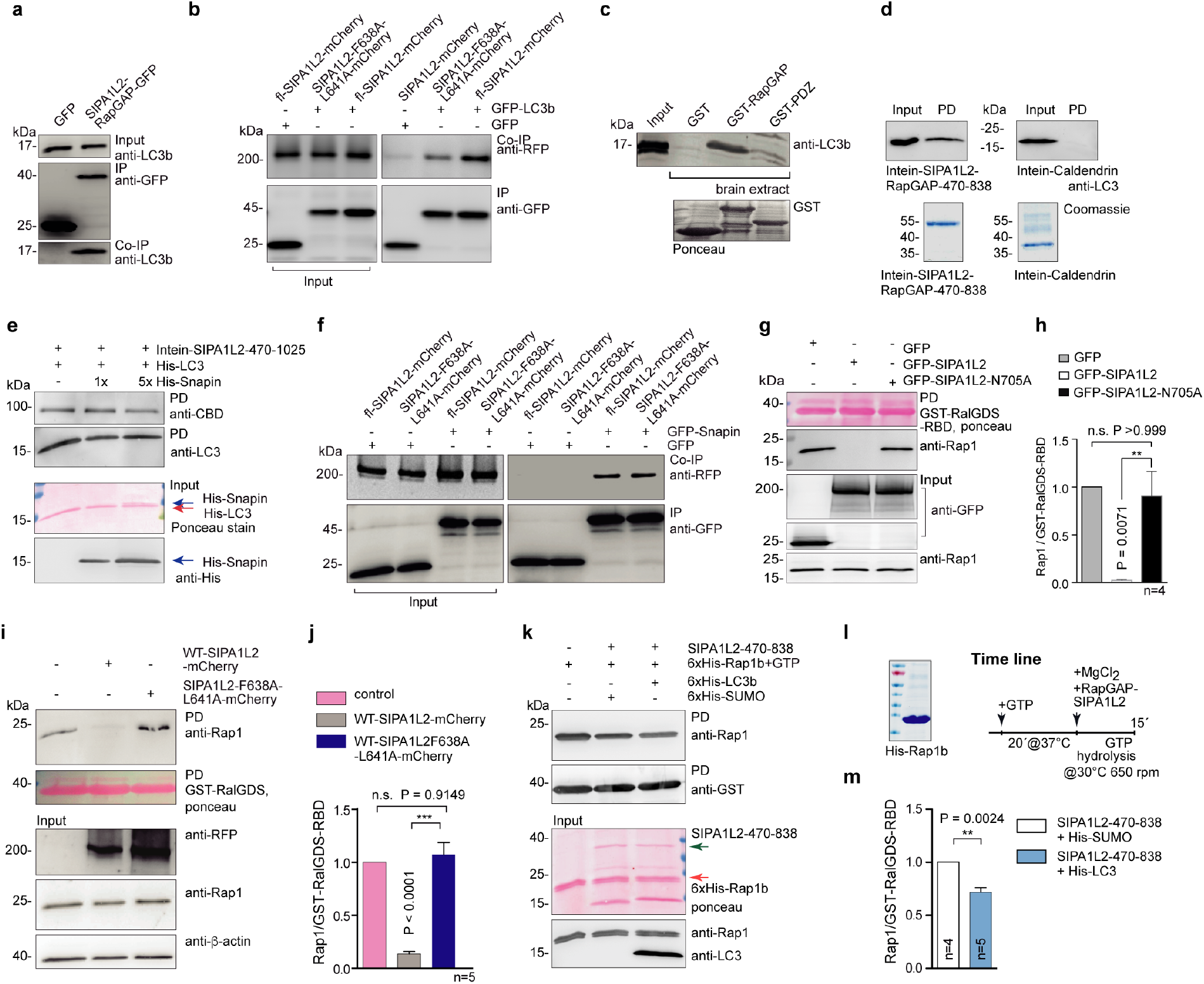
LC3 interacts with the RapGAP domain of SIPA1L2 and promotes RapGAP activity. **a.** GFP-RapGAP but not GFP alone immunoprecipitates endogenous LC3 from HEK293T cell extract. **b**. Heterologous Co-IP from HEK293t cells showing a reduced interaction between GFP-LC3b and SIPA1L2-F638A-L641A-mCherry, which harbors a mutation within the LIR motif (FxxL/AxxA) when compared to fl-SIPA1l2-mCherry. **c.** GST-RapGAP but not GST-PDZ domain of SIPA1L2 pulled down endogenous LC3 from rat brain extracts. Ponceau staining below indicates protein loading. **d.** LC3b directly interacts with Intein-RapGAP domain of SIPA1L2 (SIPA1L2-470-838), but not with Intein-Caldendrin. Coomassie blue is shown below **e**. Excess (5X) of Snapin does not interphere with the binding of LC3 to the RapGAP domain (SIPA1L2-470-1025) in a pull down assay. **f**. Both fl-SIPA1L2-mCherry and SIPA1L2-F638A-L641A-mCherry co-immunoprecipitates with GFP-Snapin. **g,h.** Rap-GAP activity assay shows decreased pull-down of Rap1-GTP (**g**) from HEK293T cell extracts in the presence of SIPA1L2-GFP but not GSP-SIPA1L2-N705A. The bar graph shows the quantification of the Rap1/RalGDS-RBD ratio from 4 independent experiments. (One-way ANOVA with Bonferroni’s correction). **i,j.** WT**-**SIPA1L2-mCherry but not SIPA1L2-F638A-L641A-mCherry reduces Rap1-GTP pull-down (**j**) with GST-RalGDS-RBD when expressed in HEK293T cells. Quantification of the Rap1/GST-RalGDS-RBD ratio is shown in **j** (One-way ANOVA with Bonferroni’s correction). **k,l,m.** Rap-GAP activity assay performed with recombinant SIPA1L2-470-838 in the presence of recombinant His-LC3b or His-SUMO as a negative control. **l.** Time line of the RapGAP activity assay and purity of recombinant His-Rap1b as confirmed by SDS-PAGE and Coomassie-blue staining. **m.** Quantification of the Rap1-GST-RalGDS-RBD ratio shows the effect of LC3b in enhancing RapGAP activity of SIPA1L2 (paired Student’s t test).

In the next set of experiments, we determined whether binding of LC3 might modulate RapGAP activity. To this end we extracted SIPA1L2 expressed in HEK293T cells and pulled down endogenous GTP-bound Rap1 with a GST-matrix. As a negative control we used a SIPA1L2 mutant carrying a point mutation within the Asn-thumb (SIPA1L2-N705A) that renders the RapGAP domain inactive (Figure S1b / Fig. 4 g,h). RapGAP activity of SIPA1L2 was abolished when we expressed the SIPA1L2 LIR-mutant (Figure 4i,j). On the contrary, RapGAP activity was enhanced in the presence of recombinant His-LC3b (Figure 4k-m). Collectively these data suggest that the interaction of LC3b with the RapGAP domain modulates the catalytic activity of SIPA1L2.

### The RapGAP activity of SIPA1L2 controls retrograde trafficking of TrkB/LC3b

Rap signaling is necessary for sustained activation of ERK (23, 24, 25) which in turn has been shown to recruit Dynein and to promote retrograde trafficking of TrkB (26). Therefore we asked whether the RapGAP activity of SIPA1L2 might modulate trafficking. Live-imaging of rat hippocampal neurons confirmed co-trajectories of LC3b and SIPA1L2 in a predominantly retrograde direction (Fig. 5a-c). LC3b trajectories were in the majority of cases positive for SIPA1L2 (Fig. 5d). A SIPA1L2-RapGAP dead mutant that is incapable of terminating Rap-signaling also showed co-trajectories with LC3b (Fig. 5c-d). However, under these conditions the velocity and the run length of the SIPA1L2-LC3b complexes were significantly enhanced (Fig. 5b,e-g), in agreement with an increased ERK activation and recruitment of Dynein (26). Finally, live-imaging of short axonal segments (30-40 µm) where active presynaptic terminals were identified using Synaptotagmin1^Oyster650^ live-antibody labeling demonstrated that SIPA1L2/LC3b complexes stopped at single boutons (Fig. 5a,b). While no changes in the number of visited boutons per axonal segment between WT- and RapGAP-dead SIPA1L2/LC3b trajectories was observed (Fig. 5h), we found much shorter dwelling times of RapGAP-dead SIPA1L2/LC3b at presynaptic boutons (Fig. 5i-j).

**Figure 5.**
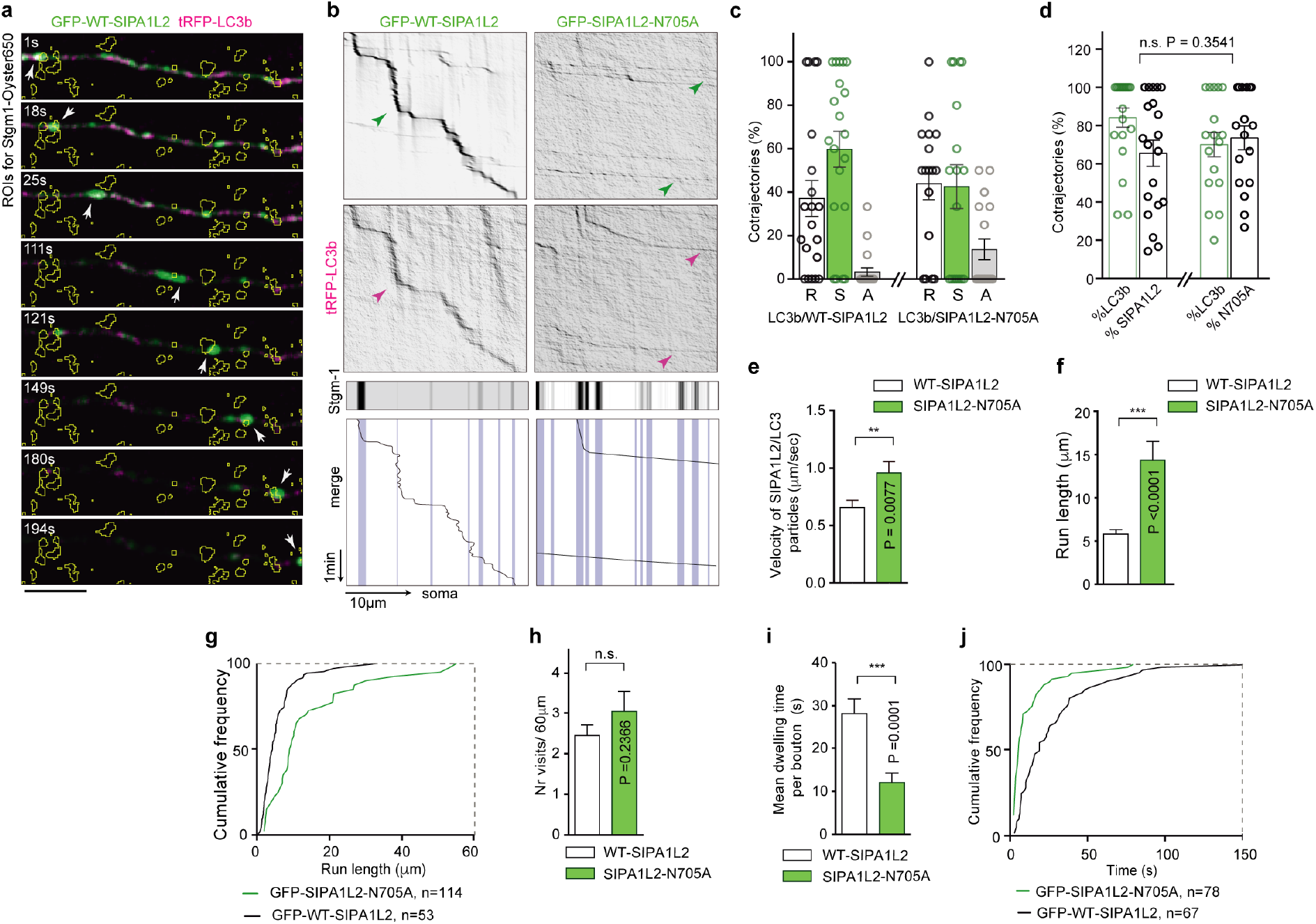
WT-SIPA1L2 co-trafficks with LC3b in axons and RapGAP activity of SIPA1L2 controls motility of the complex. **a.** Time-lapse representation of GFP-SIPA1L2/tRFP-LC3b containing organelles visiting presynaptic boutons labeled by Stgm-1^Oyster650^ (yellow ROIs). Three-channel time-lapse imaging was performed for 5 min. Scale bar = 5 µm. **b**. Representative kymographs generated from axons in (**a**). Neurons expressed tRFP-LC3b and either GFP-WT-SIPA1L2 or GFP-SIPA1L2-N705A. Below, stationary Stgm-1^Oyster650^ indicates presynaptic boutons. Sketch drawing represents traces of cotrajectories aligned with positions of presynaptic boutons (shaded blue lines). **c,d.** Percentage of WT-SIPA1L2/LC3b and SIPA1L2-N705A/LC3b cotrajectories show that both move predominantly in a retrograde (R) manner (S, stationary; A, anterograde) (**c**). Percentage of trajectories analysed per kymograph (**d**) shows a high degree of colocalizing particles and no differences between WT-SIPA1L2 and SIPA1L2-N705A. Circles in bar graphs shows values per analysed kymograph from >3 independent cultures. Bar graph shows mean±S.E.M. (one-way ANOVA with Bonferroni’s posthoc test). **e.** Velocity (μm/sec) of GFP-WT-SIPA1L2/tRFP-LC3b and GFP-SIPA1L2-N705/tRFP-LC3b cotrajectories shows that SIPA1L2 RapGAP-dead mutant moves significantly faster. WT-SIPA1L2 n=102 from 20 axons; SIPA1L2-N705A n=77 from 14 axons, >3 independent cultures. Data are depicted as mean±S.E.M (Mann-Whitney U test). **f,g.** Quantification (**f**) and cumulative distribution (**g**) of the run length (WT-SIPA1L2: n=114 from 20 axons, >3 independent cultures; SIPA1L2-N705A: n=53 from 14 axons, >3 independent cultures). **h.** Total number of visited boutons by WT-SIPA1L2/LC3b and SIPA1L2-N705A/LC3b shows no differences (Mann-Whitney U test). WT-SIPA1L2: n = 21 from 20 axons, >3 independent cultures; SIPA1L2-N705A: n=21 from 14 axons, >3 independent cultures. **i,j**. Mean synaptic dwelling time (WT-SIPA1L2: n=78 from 20 axons, >3 independent cultures; SIPA1L2-N705A: n=67 from 14 axons, >3 independent cultures). Data depicted as mean±S.E.M. Mann-Whitney U test.

### Induction of cLTP prolongs dwelling time of the SIPA1L2-TrkB-LC3 amphisome at presynaptic boutons

Presynaptic LTP results in activation of Protein Kinase A (PKA) at MF boutons and enhanced synaptic function that is largely mediated by PKA-dependent phosphorylation of different components of the presynaptic release machinery (27, 28, 29, 30, 31). PKA-mediated phosphorylation of Snapin at Ser50 is crucial for its dissociation from the DIC and a phosphomimetic Snapin-S50D mutant is largely immobile (32). Heterologous co-immunoprecipitation experiments revealed that both, phosphomimetic (S50D) as well as phosphodeficient (S50A) Snapin interact with tRFP tagged SIPA1L2-470-1025 (Fig. 6a). However, in contrast to the phospho-deficient protein, DIC did not co-immunoprecipitate with phosphomimetic Snapin (Fig. 6a), indicating that PKA-dependent phosphorylation induces the dissociation of the protein from DIC. Consistent with previous work (32), time-lapse imaging revealed that phosphomimetic Snapin is largely stationary and was often found in proximity to immobile WT-SIPA1L2-mCherry at axon terminals (Fig. 6b, c). We therefore next wondered whether changes in synaptic activity could affect amphisome trafficking among presynaptic terminals. Trafficking of SIPA1L2/LC3 among presynaptic terminals labeled by anti-Synaptotagmin1 antibodies revealed an enhancement in dwelling time upon induction of cLTP induction (Fig. 6d-f). The PKA inhibitor H89 prevented this effect (Fig. 6e,f).

**Figure 6.**
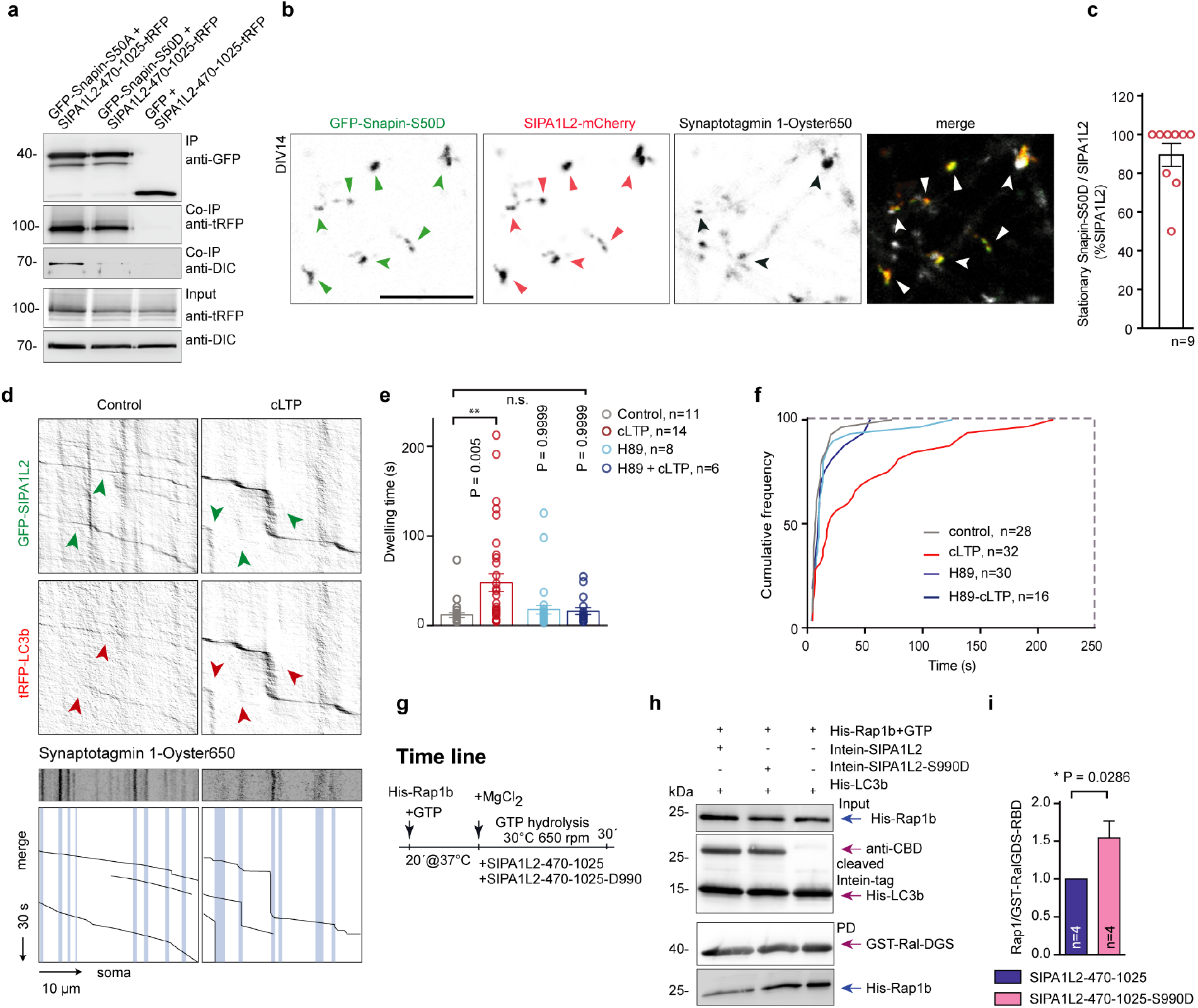
cLTP prolongs dwelling time of the SIPA1L2-containing amphisome at presynaptic boutons. **a.** Heterologous co-immunoprecipitation experiments showing that phosphodeficient (GFP-Snapin-S50A) but not phosphomimetic Snapin mutant (GFP-Snapin-S50D) co-immunoprecipitates with DIC. **b,c.** Snap-shots obtained from time-lapse imaging in cells overexpressing GFP-Snapin-S50D together with WT-SIPA1L2-mCherry where mostly immobile (arrows) vesicles were found at presynaptic terminals labelled with anti-Synaptotagmin 1-Oyster650 (arrows). Scale bar is 10 µm. In (**c**) quantification of the percentage of stationary co-trajectories. **d,e,f.** Kymographs generated from axons co-expressing GFP-SIPA1L2 and tRFP-LC3b and labeled *in-vivo* with Stgm-1-Oyster650 after control or cLTP induction. In the bottom, merge images represent traces of co-trajectories aligned with positions of presynaptic boutons (shaded blue lines). Dwelling time of the SIPA1L2-LC3b-amphisomes per bouton is represented in (**e**) and corresponding cumulative distribution diagram in (**f**). Data represented as mean±S.E.M. (One way ANOVA on ranks with Dunn’s multiple comparison test). **g,h,i.** RapGAP activity assays show a negative regulation of SIPA1L2 catalytic activity by potential PKA phosphosite (Ser990). Recombinant intein-tagged SIPA1L2-470-1025 as well as SIPA1L2-470-1025-S990D were used to hydrolize recombinant Rap1b loaded with GTP in the presence of LC3b. Quantification of 4 independents experiments in **i** (Mann-Whitney U test).

It has been reported that PKA-dependent phosphorylation of Ser499 of Rap1GAP negatively regulates RapGAP activity (33). Sequence analysis of SIPA1L2 revealed a high scoring PKA phosphorylation motif (S990-RxxpS motif/ 0,757 / Fig. S1d) that is located close to the region that corresponds to the regulatory PKA sites (pSer 499) of Rap1GAP and that is well preserved between SIPA1L family members (Fig. S1b). When we tested the hypothesis that PKA in analogy to Rap1GAP might negatively regulate RapGAP activity of SIPA1L2 we found that phosphomimetic SIPA1L2-470-1025-S990D indeed hydrolized very little recombinant Rap1b-GTP (Fig. 6g-i), indicating a negative regulation of RapGAP activity by PKA. Collectively these data suggest that following induction of presynaptic plasticity, PKA phosphorylation of Snapin induces the dissociation of the amphisome from Dynein and enhances its residing time at presynaptic boutons. Concomitant phosphorylation of SIPA1L2 diminishes its RapGAP activity and thereby facilitates ERK signaling.

### TrkB associates at presynaptic sites to amphisomes

Amphisomes are transient intermediate organelles formed upon the fusion between autophagosomes and endosomes that in non-neuronal cells rapidly enter a degradative lysosomal pathway. However, lysosomes are not abundant if at all present in distal axons (34). In primary cultures, STED imaging revealed that the lysosomal marker LAMP1 was indeed mostly absent from presynaptic boutons, making it unlikely that SIPA1L2-TrkB-LC3b associates with a degradative organelle at axon terminals (Fig. S5k). A recent study showed that TrkB is transported on autophagosomes to the soma where it regulates gene expression (35). This transport requires association of the Clathrin adaptor AP2 with LC3 and DIC (35). We therefore speculated that amphisomes could constitute more persistent entities in axons where they could serve signalling functions. Accordingly, we found that SIPA1L2/LC3/TrkB complexes are positive for the late endosome marker Rab7 (Fig. 7a). Moreover, kinase-active pTrkB^Y515^ showed a significant degree of colocalization with SIPA1L2 at presynaptic boutons in line with the high association of LC3, SIPA1L2 and TrkB observed before (Fig. 7b-c; Fig. S5g,h). This is expected to be higher when assessing colocalization in only those boutons in which SIPA1L2 is present. Accordingly, intensities of pTrkB^Y515^ are higher in SIPA1L2-positive boutons (Fig. 7d-e). The tight association of SIPA1L2 with pTrkB^Y515^ (Fig. 7b-c) as well as TrkB with LC3 (Fig. 7f,g) and Rap1 with SIPA1L2 and LC3 (Fig. S5l) at presynapses indicates that SIPA1L2 might be part of a signaling amphisome. Manders’ coefficient also shows extensive colocalization with AP2 in axons and synaptic boutons (Fig. S6a-c), suggesting that SIPA1L2-TrkB amphisomes are long-range organelles that will traffic retrogradely to reach the soma (35). Moreover, TrkB and SIPA1L2 associate with an autophagosome enriched fraction from rat brain lysates (Fig. 7h,i) that is also positive for LC3, Rab7 and Snapin. In addition, we could identify TrkB tagged at the C-terminus with GFP as part of double membrane organelles in the MF projection with immunogold EM following low-level expression of the fusion protein in organotypic slices. Staining with a GFP-antibody resulted in immunogold particles localized to the outer membrane (Fig. S6d) in agreement with a potential signaling function of the organelle.

**Figure 7.**
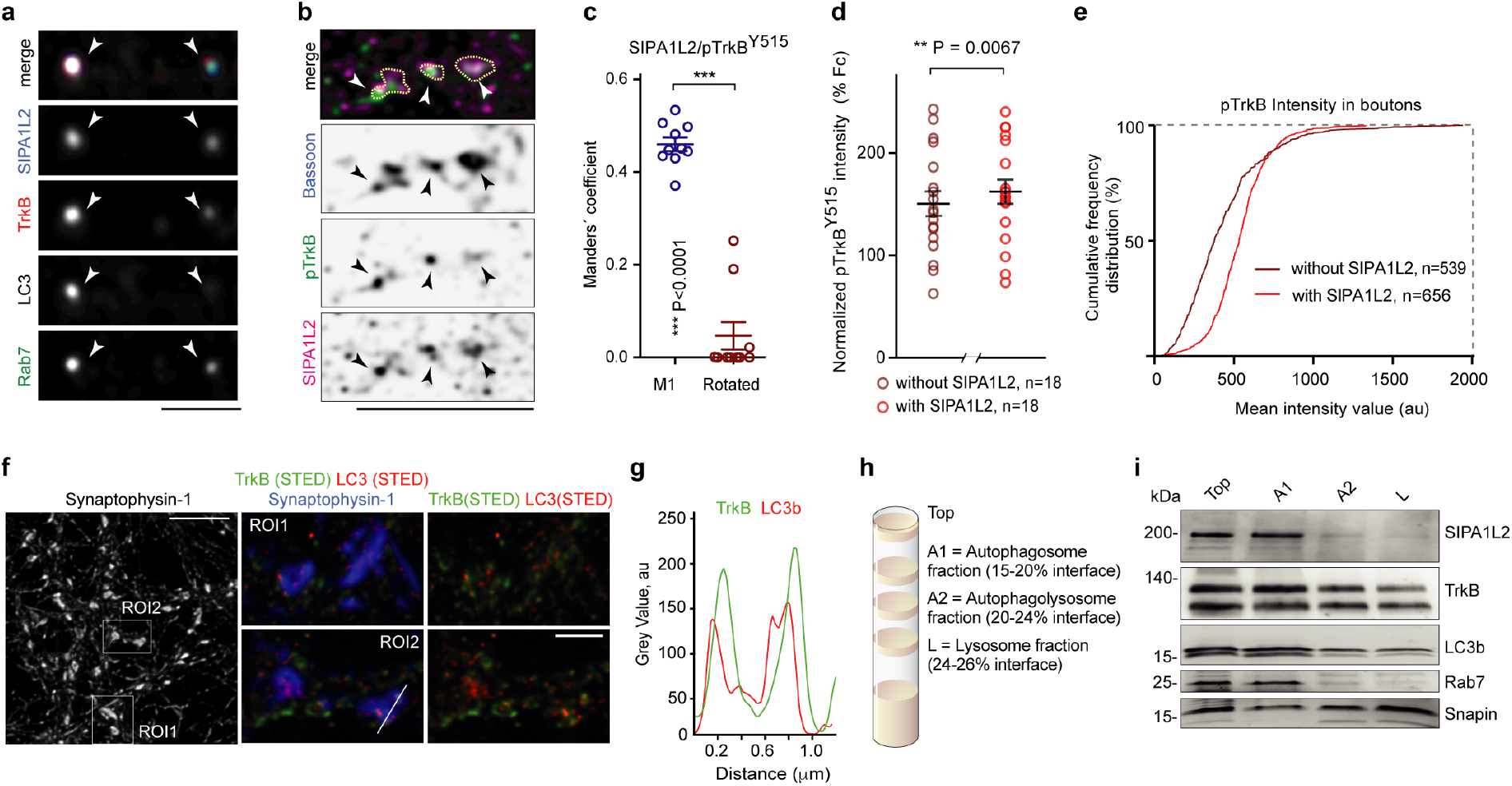
TrkB-LC3b-Rab7 containing amphisomes are positive for SIPA1L2. **a.** Confocal images of a quadruple immunofluorescence performed in rat primary hippocampal neurons against SIPA1L2, LC3, TrkB with Rab7. Scale bar is 1 µm. **b**. Confocal images of primary neurons treated with BDNF and stained for pTrkB ^Y515^, SIPA1L2 and Bassoon show colocalization of SIPA1L2 and pTrkB ^Y515^ in presynaptic boutons. Scale bar is 5 µm. **c.** Mander’s coefficient calculated for pTrkB^Y515^ and SIPA1L2 in boutons detected by Bassoon staining. As control, same images were rotated 90° to the right. Paired Student’s t test. **d,e.** pTrkB^Y515^ intensities after BDNF treatment and calculated in boutons labeled by Sphy 1 and segregated by the presence or absence of SIPA1L. Data was normalized to images acquired in neurons treated with Trk-Fc and represented as averaged intensities for boutons per image (d, “n” - indicates number of images), as well as cumulative frequency distribution (e, “n” - indicates number of synaptic boutons). Paired Student’s t test. **f,g.** Super-resolution STED imaging revealed association of TrkB with LC3 at the presynaptic boutons. Scale bars 5 µm (overview) and 1 µm (inserts). **h,i.** SIPA1L2, TrkB, Rab7 and Snapin are present in an autophagosome enriched fraction following subcellular fractionation (scheme in **h**).

What could be the consequence of the amphisome stopover at axon terminals? Local ERK activation has been linked to enhanced neurotransmitter release and presynaptic plasticity (36, 37). We took advantage of a FRET-based ERK sensor (38) that we targeted to the presynaptic compartment following fusion to Synaptophysin-1 (Fig. 8a, S6e). We then imaged changes in the GFP fluorescence lifetime, the donor of the FRET pair, coinciding with the stopover of amphisomes, which we identified using either tagged versions of SIPA1L2 or LC3. Synaptic visits were determined by kymograph analysis (Fig. 8a,b). Quantification of ΔLifetime_GFP_ upon the arrival of the amphisome (Fig. 8c,d) at the bouton (time 0) indeed correlated with a significant decrease in the GFP lifetime (Fig. 8b-e), indicating a higher FRET efficiency due to ERK activation and subsequent phosphorylation of the sensor (38). Importantly, this decrease was not observed when the amphisome passed by a bouton without stopping (non-visited terminals) or in axons where no amphisomal trafficking was observed during the recording time (Fig. 8e).

**Figure 8.**
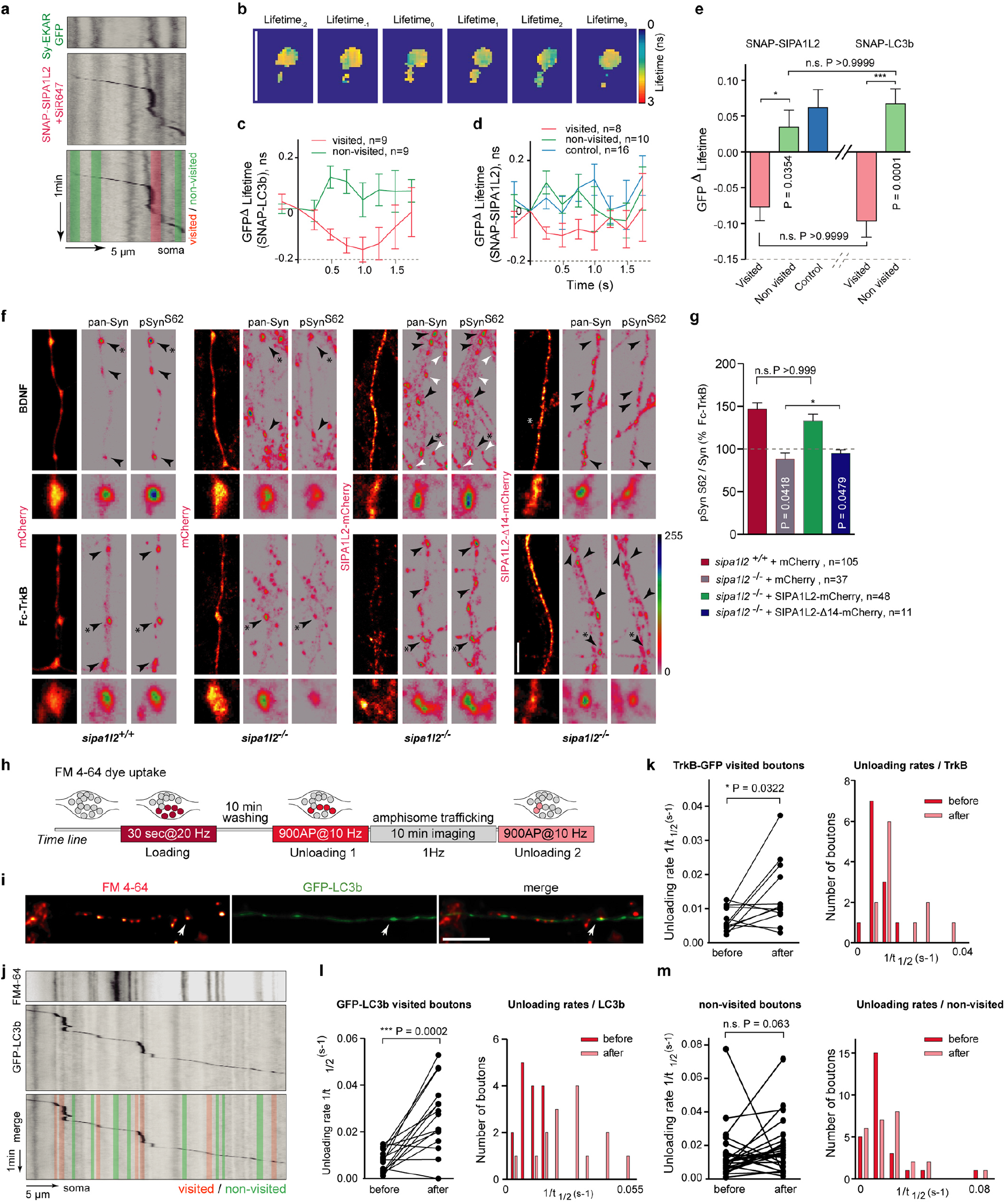
SIPA1L2 amphisomes activate ERK at presynaptic boutons and potentiate presynaptic function. **a.** Kymographs represent the visit (red-shaded line) of SIPA1L2 containing amphisome labeled by SNAP-SIPA1L2 (+SiR647) to a bouton identified by Sy-EKAR. Non-visited boutons are marked with a green shaded line. **b.** Representative heat maps depicting GFP lifetime (ns) over time. Lifetime _0_ represents the frame when the amphisome reaches the bouton. Scale bar is 5 µm. **c,d.** Quantification of GFP-ΔLifetime over time in visited and non-visited boutons. Arrival of the amphisome is considered as time_0_. Control in (**d)** represents data from axons where no trajectories were found. N numbers represent analysed boutons from n>3 independent experiments. **e.** Averaged GFP-ΔLifetimes in **c,d** after time 0. Bars represent mean±S.E.M. Amphisomes are detected as SNAP-LC3b (+SiR647) or SNAP-SIPA1L2(+SiR647)-positive organelles. Number of boutons are shown in **c,d**. (One-way ANOVA with Bonferroni’s posthoc test). **f.** Representative confocal images from wt and *sipa1l2*^−/−^ primary neurons overexpressing WT-SIPA1L2-mCherry, SIPA1L2-Δ14-mCherry or mCherry as control, treated with either Fc-TrkB or BDNF and immunostained for Synapsin1,2 (Syn) and phospho-Syn^S62^. White arrows show non-transfected boutons. Insets are zoomed in from boutons labeled with the asterisk. Scale bar is 5µm. **g.** Percentage of enhancement in the pSyn^S62^/Syn ratio in BDNF-treated as compared to Trk-Fc treated neurons). Data shown as mean±S.E.M. Number of analysed boutons are in the legend. (One-way ANOVA with Bonferroni posthoc test). **h.** Schematic representation showing the time-line of the experiment: FM_4-64_ (10µM) was loaded in the terminals by a train of pulses (30 sec @ 20Hz) and washed out for 10 minutes. First FM_4-64_ unloading was done by delivering 900 pulses@10Hz. Trafficking of TrkB-GFP or GFP-LC3b was imaged for 10 min before a second unloading protocol was applied. **i.** Representative images showing an axon overexpressing GFP-LC3b after loading with FM4-64 (red). Scale bar is10 µm. **j.** Kymographs prepared from an axonal segment overexpressing GFP-LC3b and loaded with FM4-64. Below, a schematic representation shows the discrimination made between visited (red) and non-visited (green) boutons. **k,l.m.** Quantification of the unloading rates obtained in axons overexpressing TrkB-GFP (**k**) or GFP-LC3b (**l**) and non-visited boutons as control (**m**). Each dot represents an analysed bouton. Right panels show the frequency distribution of unloading rates. (Paired Student’s t-test in **k**, **l** and Wilcoxon rank test in **m**).

ERK activation in presynaptic plasticity results in phosphorylation of Synapsin I (Syn-I), a protein, which is known to play a role in BDNF-mediated increase of neurotransmitter release (37). Ultrastructural analysis revealed no major alterations in presynaptic bouton organization and vesicle content in *sipa1l2*^−/−^ as compared to *wt* mice (Fig. S6f,g). Quantification of ERK-dependent phosphorylation of Syn-I in hippocampal primary neurons from *wt* mice revealed a significant enhancement of the pSyn/Syn ratio at boutons following BDNF application (Fig. 8f,g) without changes in their number (Fig. S6h). The BDNF-induced enhancement of the pSyn/Syn ratio was absent in *sipa1l2*^−/−^ neurons and could be rescued by re-expression of WT-SIPA1L2 in ko neurons. However, no rescue was observed when we re-expressed the SIPA1L2-Δ14 mutant lacking the TrkB binding region (Fig. 8f,g).

To assess transmitter release directly we combined unloading of FM 4-64 dyes to monitor synaptic vesicle fusion with live imaging of LC3b/TrkB trafficking (Fig. 8h,i) and compared unloading rates at visited and non-visited boutons (Fig. 8i,j). Analysis of the unloading rates before and after the stopovers revealed a significant enhancement in the majority of the analyzed boutons visited by both TrkB-GFP (Fig. 8k) and GFP-LC3b (Fig. 8l). This increase was not observed in non-visited neighbouring boutons (Fig. 8m).

## Discussion

Autophagosomes in neurons are formed continuously in distal axon terminals (39) from where the autophagic flux of cargoes derived from synapses is directed towards the soma in a Dynein-dependent manner (40, 39, 41, 42, 43, 20). Classical stimuli like starvation have relatively little effect on neuronal autophagic flux (41). However, it was shown previously that synaptic activity transiently up-regulates autophagy at presynaptic terminals (44, 45, 46, 42) and autophagosome formation might be therefore regulated locally in individual boutons. Although there is growing appreciation that neuronal autophagy might serve synaptic function (47, 48, 49) it has not been shown yet that it might be a mechanism used to mediate activity-dependent synaptic change. In the present work we introduce a signaling mechanism that is based on incorporation of TrkB receptors from late endosomes to autophagosomes. This fusion has been largely associated with the termination of signaling (50; 51) and evidence for the existence of a signaling amphisome was scarce (52). We found that the resulting organelle has indeed features of an amphisome with signaling properties whose trafficking and signaling capabilities are tightly controlled by SIPA1L2 and its association with TrkB, Snapin and LC3. Intriguingly, binding of LC3 enhances RapGAP activity and thereby negatively interferes with TrkB-induced Rap signaling, which is upstream of ERK activation (Fig. S7). PKA phosphorylation of Snapin and SIPA1L2 at presynaptic sites immobilizes the amphisome at axon terminals, terminates RapGAP activity and thereby allows TrkB/Rap1 signaling that will facilitate transmitter release (Fig. S7). It was proposed previously that autophagosomes contain active TrkB complexes (35). In this study we provide evidence that TrkB signaling endosomes are in fact amphisomes whose formation is likely linked to the regulation of neuronal autophagy. We propose that neuronal amphisomes are stable prelysosomal hybrid organelles based on the findings that i) they were immunopositive for the late endosome marker Rab7 but not the lysosomal marker LAMP1, ii) TrkB-GFP is localized to the outer membrane of autophagosomes in MF axons, iii) in presynaptic boutons TrkB is invariably found in close proximity to SIPA1L2 and phosphorylated at Y515, which is a crucial phosphosite for ERK-signaling, iv) amphisome trafficking to boutons correlated with enhanced levels of phosphorylated Syn-I and ERK activation and v) finally, enhanced transmitter release.

What could be the advantage of assembling a stable hybrid organelle for signaling at axonal boutons? Axon terminals and distal axons contain very few if any lysosomes and acidification and proteolytical cleavage is significantly delayed until amphisomes reach the soma (34, 40). Most important, whereas the lumen of autophagosomes becomes more acidified during retrograde transport and lysosomal markers accumulate, levels of cathepsins, the major lysosomal proteases, remain very low compared to those in the soma (53). If one takes into account the extreme distances in axonal transport, amphisomes will constitute stable entities long before acquiring a lysosomal identity near the neuronal cell body. Since autophagosomes fuse with late endosomes in order to undergo robust retrograde transport (20) it is probably inevitable that, in the absence of autolysosome formation, amphisomes serve as signaling and sorting platforms while trafficking in a retrograde direction. This makes endosomal sorting processes at axon terminals dispensable and would give an answer to the long-standing question of why neurons transport autophagic and endocytic cargos back to the cell body for degradation instead of disposing them locally.

It is likely that amphisomes will collect additional cargo during visits at synapses that may differently determine their properties and vesicular fate while propagating retrogradely to the soma. Sustained signaling capacity and diversity will promote and integrate local synaptic function with long-range signaling while evading lysosomal degradation. Specificity for certain synaptic connections is conceivable and in the case of SIPA1L2 might come from docking to presynaptic sites via for instance PDZ domain-dependent interactions. The idea of selective amphisome signaling at subpopulations of boutons is appealing since TrkB/BDNF facilitation of presynaptic transmitter release possibly requires more sophisticated signaling than the one provided by the classical TrkB signaling endosomes that lack local signaling function and selectivity for individual boutons required to induce clustered presynaptic plasticity like it was proposed by Staras et al. (54). This study demonstrated the existence of a vesicle pool that is not confined to a synapse but spans multiple terminals. Vesicles within this superpool are highly mobile and are rapidly exchanged between terminals. Focal BDNF application suggests the involvement of a local TrkB-receptor-dependent mechanism for synapse-specific regulation of presynaptic vesicle pools through control of vesicle release and capture to or from the extrasynaptic pool (54). Thus, clustered presynaptic plasticity is another conceivable function of amphisome signaling. This is also interesting in light of several studies which have shown that MF LTP is expressed presynaptically and requires cAMP–PKA signaling (55). Our results suggest that the SIPA1L2 amphisome is a likely substrate for PKA-dependent LTP. Along these lines we speculate that at MF synapses SIPA1L2 related amphisome signaling is important for the maintenance of LTP, which in turn requires BDNF/TrkB signaling and is important for pattern separation.

Several adaptors have been described for antero- and retrograde transport of TrkB (56, 19, 57, 58, 59, 35). Many of these adaptors show also an association with authophagosomes (60, 61, 62) and it is likely that synergies exist between adaptors. Interestingly, Snapin is reportedly also an adaptor for axonal lysosomes (63), which further supports the idea of a single transport complex slowly acquiring a lysosomal identity during retrograde movement. We could show that SIPA1L2 mediated ride-on service for TrkB-containing amphisomes enables local and long-distance signaling (Fig. S7). In this context, it should be mentioned that other SIPA1L family members exhibit a similar domain organization with a high degree of similarity in the RapGAP, PDZ and CC-leucine zipper domain (Fig. S1). It is thus very likely that all will bind Snapin and LC3 while they might differ in cargo selection for retrograde transport of amphisomes but also additional functions of the SIPA1L family in autophagy are conceivable.

## SUPPLEMENTAL INFORMATION

Supplemental Information includes seven figures can be found with this article online at …

## Acknowledgements

The authors gratefully acknowledge the professional technical assistance of C. Borutzki, S. Hochmuth and M. Marunde, O. Kobler for help with imaging data processing, M. Lever for help with cloning. Supported by grants from the Deutsche Forschungsgemeinschaft (DFG) (KR 1879 / 9-1 / FOR 2419, Kr1879 / 5-1 / 6-1 / SFB 779 TPB8), BMBF ‘Energi’ FKZ: 01GQ1421B, The EU Joint Programme – Neurodegenerative Disease Research (JPND) project STAD and Leibniz Foundation to MRK; DFG SFB 779 TPB8 to AK; DFG STO488/4-1 / SFB779 TPB5 to OS; DFG 556/11-1 to MK. Leibniz SAW to EDG; EU QLG3-CT-2001-01181 to MC, EDG. GMG was supported by a Alexander-von-Humboldt Foundation / CAPES post-doctoral research fellowship.

## Author information

### Contributions

M.A.-A., C.S., A.K., M.R.K. designed the study, M.A.-A., A.K., M.R.A., I.B., G.A.S., G.M.G., R.R., P.Y., M.B., J.L.-R., T.H., T.M., C.S. conducted the experiments, M.A.-A., A.K., M.R.A., I.B., G.A.S., G.M.G., P.Y., M.B., J.L.-R., S.D.-G., M.S., T.M., C.S., M.R.K. analyzed data, R.R., O.S., N.B., M.C., A.V.F., A.R., M.S., M.K., E.D.G., provided tools and reagents, M.A.-A., A.K. and M.R.K. wrote the paper and all authors commented and revised the manuscript.

## Supplemental Information

**Figure S1.**
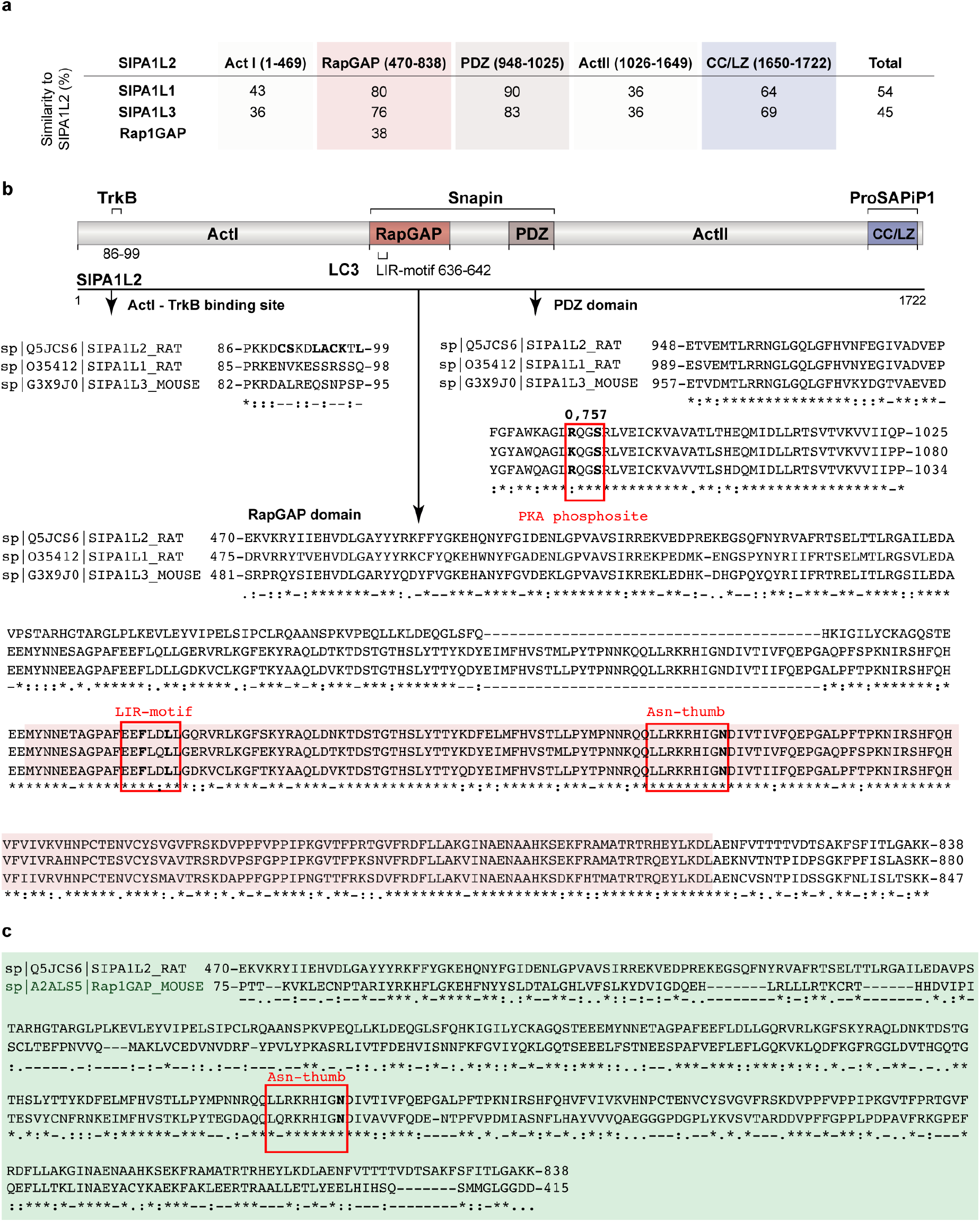
SIPA1L2 shows a conserved domain organization and a high degree of similarity with other SIPA1L family members and Rap1GAP. **a.** Table showing the similarity of SIPA1L2 with SIPA1L1 and SIPA1L3 as well as the RapGAP domain of Rap1GAP. Comparisons were made according to the SIPA1L2 domains defined by the amino acids depicted in the heading. The RapGAP and PDZ domains are highly conserved among SIPA1L family members. **b.** Scheme showing the domain organization and interaction motifs of SIPA1L2 with TrkB, Snapin, LC3 and ProSAPiP1 (12). Depicted the sequence comparison of the ActI, RapGAP and PDZ domains between SIPA1L family members. Boxes within the RapGAP domain indicate the LIR-motif and the Asparagine thumb where amino acids in the so-called LIR-mutant (F638A-L641A) and RapGAP dead mutant (N705A) are indicated in bold. Otherwise stated, interaction assays performed in this study involving the RapGAP or PDZ domain were done with SIPA1L2-624-813 (SIPA1L2-RapGAP; red shaded box in the figure) and SIPA1L2-948-1025 (SIPA1L2-PDZ). The alignment for the RapGAP domain is done with a longer sequence stretch (470-838) defined by comparison with a published, catalytically-active fragment of Rap1GAP (aa 75-415; 23). Fragment SIPA1L2-470-838 was used in this study in RapGAP activity assays. Symbols represent: fully conserved residues (*), groups with strongly similar properties (:) and groups with weakly similar properties (.) according to ClustalW2. **c.** Sequence comparison of SIPA1L2 with the catalytically active RapGAP domain of Rap1GAP. Symbols represent: fully conserved residues (*), groups with strongly similar properties (:) and groups with weakly similar properties (.) according to ClustalW2.

**Figure S2.**
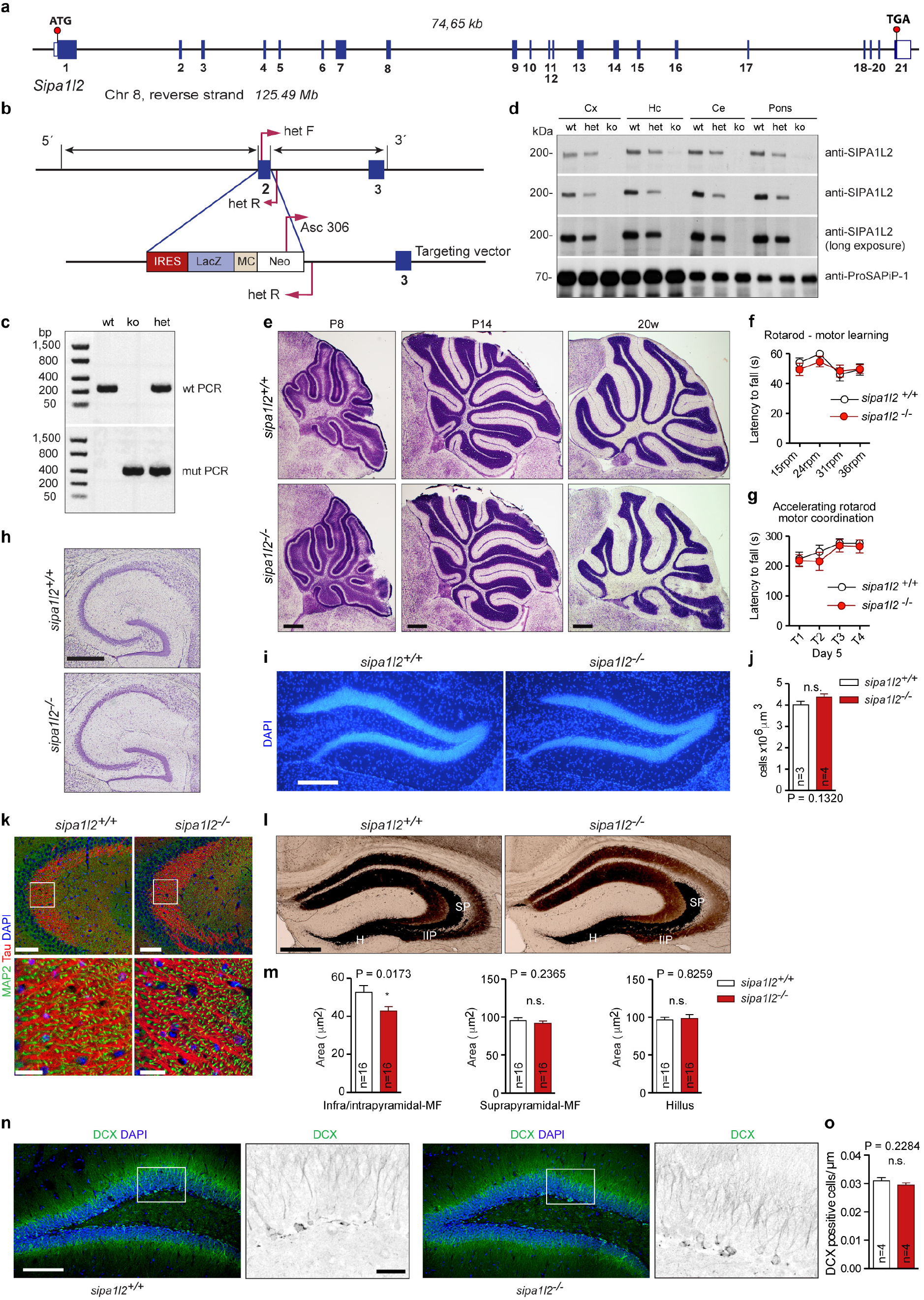
Generation and characterization of *sipa1l2* knockout mice. **a.** Schematic representation of the SIPA1l2 genomic structure comprising 21 exons located on mouse chromosome 8 with one alternatively spliced exon (no. 18). Blue boxes represent exons. **b.** Scheme depicting the strategy used for generation of the constitutive *sipa1l2* knockout mice. The targeting vector replaces exon 2 with an IRES-LacZ sequence. The positions of primers for genotyping analysis (hefF, hetR and Asc306) are indicated by pink arrows. **c.** Representative gel of mouse genotyping by PCR with primers indicated in b. Amplification with primers hetF and hetR (*wt* allele) yielded a 200-bp product, whereas amplification with primers Asc306 and hetR (recombinant allele) yielded a 400-bp product. **d.** Western blots on mouse brain tissue from different areas of *wt*, SIPA1l2 het and ko mice (Cx, Cortex, Hc, hippocampus, CE, cerebellum, Pons). Specific SIPA1l2 antibodies from different species (guinea pig and rabbit) were used to show that SIPA1l2 protein was reduced in heterozygous mice and not present in knock out animals. The expression of a SIPA1l2 interaction partner (ProSAPiP1) was not changed. **e,f,g.** *sipa1l2*^−/−^ mice display no differences from wt animals in cerebellar organization and no alteration in rotarod-motor learning and accelerating rotarod motor coordination. Nissl stained saggital cerebellar sections (**e**) from *sipa1l2*^−/−^ in comparison to *sipa1l2*^+/+^ mice show no anatomical abnormalities in the cerebellum at different developmental stages. Scale bar = 500 µm. In **f**, *sipa1l2*^−/−^ and *sipa1l2*^+/+^ mice were submitted to 4 days of rotarod task with increased speeds each day (maximum latency of 60 sec). (**g**) On day 5 mice were submitted to the accelerating rotarod task 4 times (maximum latency 300 sec). Data are expressed as the mean + S.E.M. (ANOVA), n=9. **h,i,j.** Nissl staining reveals no alterations in the hippocampus (**h**) (scale bar = 500 µm). DAPI staining in the DG reveals no alteration in the cell number between genotypes (**i,j**) (scale bar =150 µm). **k,l,m.** *sipa1l2*^−/−^ mice display minor alterations of the Mossy Fiber projection compare to wild type animals as depicted with Tau and Timm staining. (**l**) Images showing TIMM-possitive structures in hippocampus. In (**m**), H stands for Hilus, SP for suprapyramidal mossy fibers and IIP for infra-and intrapyramidal mossy fibers. Data is presented as means + S.E.M. (Mann Whitney U test for infra/intrapyramidal and suprapyramidal regions and Student’s t test for quantification in the Hillus). Scale bar =100 μm and in zoom in ROIs scale bar is 20 μm. Scale bar for Timm staining is 400 μm. **n,o.** No reduction in the number of doublecortin (DCX) positive adult-born granule cells observed in *sipa1l2*^−/−^ mice. Scale bar =100 µm and 20 µm inside the ROIs.

**Figure S3.**
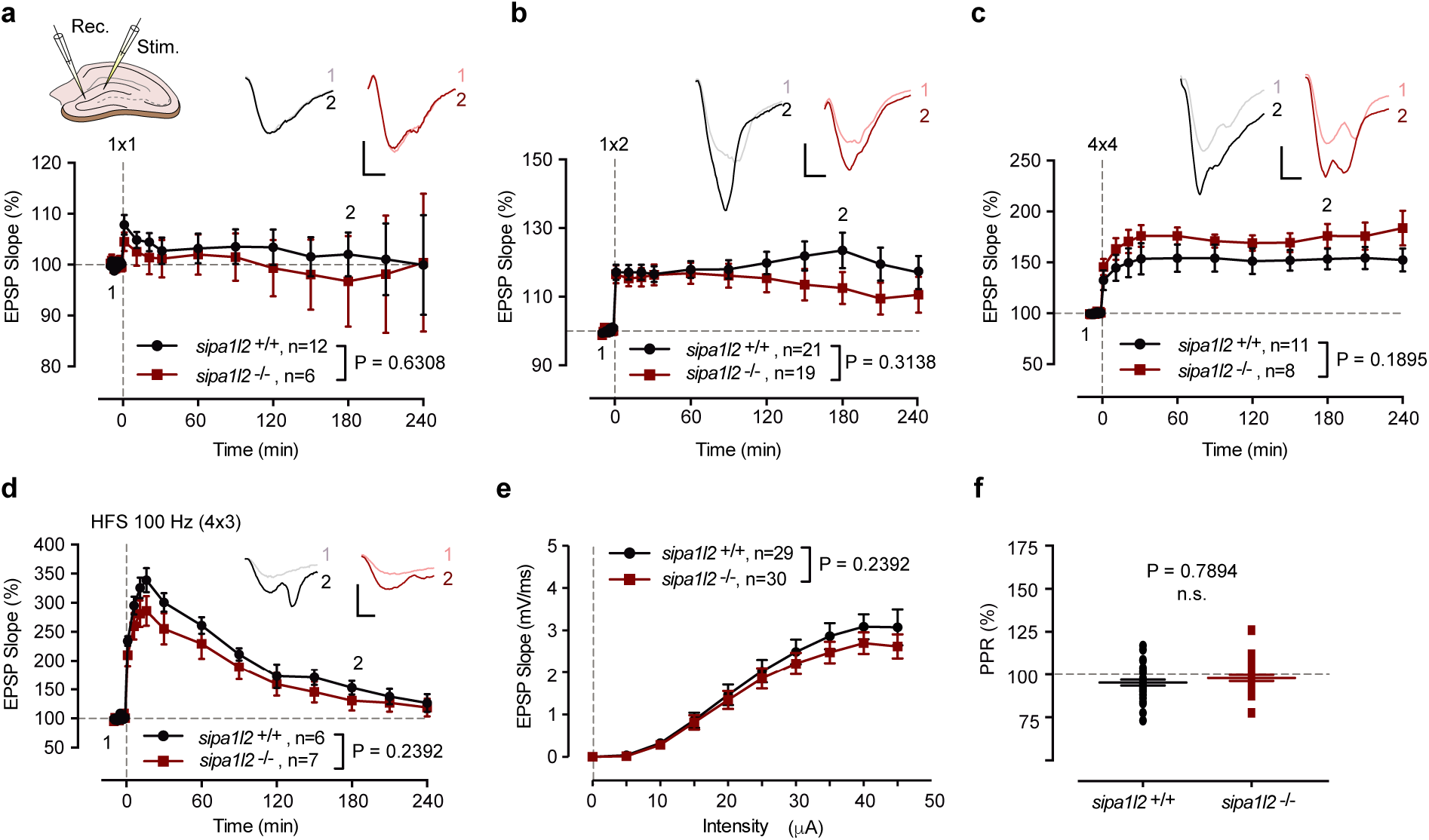
Basal synaptic transmission and synaptic plasticty in the DG of *sipa1l2*^−/−^ animals was similar to *wt* mice. **a,b.** Theta burst stimulation protocols (TBS) of different strengths delivered on acute slices from wt and *sipa1l2*^−/−^ mice elicited similar potentiation in both genotypes. Delivery of one (**a**) brief presynaptic burst of 10 pulses at 10 Hz - 0.2ms per half-wave duration - repeated 10 times at 5Hz episode of (**a**) or two of these episodes (inter-episode interval 10s) (**b**) produced similar responses in both experimental groups (two-way ANOVA). **c-d.** The potentiation levels after a stronger TBS protocol consisting of 4 episodes (inter-episode interval 10s) repeated 4 times every 5min (**c**) or a high-frequency stimulation protocol (4×100Hz 1s, every 5min) (**d**) did not differ among the groups (two-way ANOVA). **e-f.** No differences in input-output curve of the field excitatory postsynaptic potential (**e**) and paired-pulse facilitation (**f**) between *sipa1l2*^+/+^ and *sipa1l2*^−/−^ were observed (Student’s t test).

**Figure S4.**
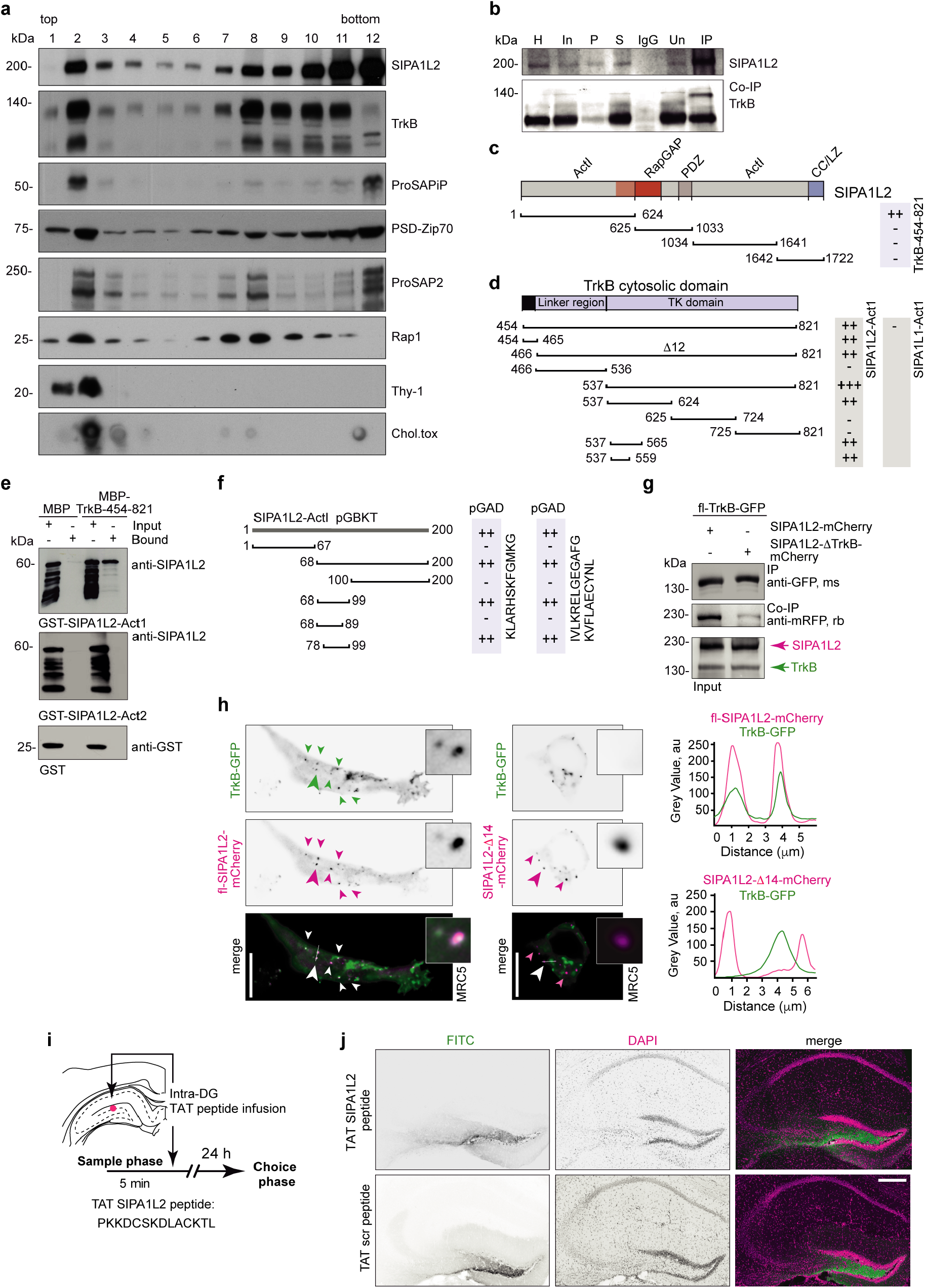
SIPA1L2 associates with TrkB. **a.** Lipid raft isolation from rat cerebellum shows that SIPA1L2 distribution overlaps with TrkB and Rap1 and associates with lipid rafts identified by Thy-1. As control, fractions were also assayed by sphingolipids-dot blot assay using Cholera toxin. P12 cerebellum from rat was extracted in 3% Brij 35 and separated on a density gradient formed from 40, 30 and 5% sucrose. Twelve fractions (from top to bottom of the gradient) were immunoblotted for indicated proteins. **b.** TrkB co-immunoprecipitates with endogenous SIPA1L2 from rat cerebellar extracts. SIPA1L2 was immunoprecipitated with anti-SIPA1L2 guinea-pig antibodies (IP) and guinea pig IgG was used as a control. Western blot of precipitates was detected with anti-SIPA1L2 rabbit and with anti-TrkB mouse antibodies. **c.** Schematic representation of SIPA1L2 domains and the corresponding fragments used in YTH (black lines) to map the binding region in SIPA1L2 to cytosolic TrkB (TrkB-454-821). Results are depicted in the light-blue box on the right that shows that the ActI domain of SIPA1L2 interacts with the intracellular domain of the TrkB receptor. Numbers indicate amino acid (aa) residue position in each protein according to the rat sequence. **d.** Scheme depicting the fragments used in YTH and their corresponding interactions between ActI-SIPA1L2 and the cytoplasmic domain of TrkB. Two binding interfaces in TrkB were found to interact with ActI-SIPA1L2: 12aa within the juxtamembranous region of TrkB (TrkB-454-465) and the first 23aa the tyrosine kinase (TK) domain (TrkB-538-560). No interaction was found between cytosolic TrkB and ActI-SIPA1L1. **e.** MBP-pull-down assays confirmed the direct interaction of ActI domain of SIPA1L2 with the cytosolic region of TrkB (MBP-TrkB-454-821) but not to MBP control. A band at about 60kDa is visualized in the WB using anti-SIPA1L2 (rb) antibodies. The ActII domain of SIPA1L2 (68 kDa; GST-SIPA1L2-1026-1650) as well as GST alone does not interact with cytosolic part of TrkB. GST-Act1-SIPA1L2-1-624 was detected with rabbit anti-SIPA1L2 (75-3); Buffer composition: TRIS 20 mM pH8.0, DTT 1 mM, EDTA 3 mM, NaCl 100 mM, Tween-20 0,3%, complete protease inhibitor. **f.** Identification of the TrkB-binding interface within the Act1 domain of SIPA1L2 by YTH. First 200 aa of the ActI domain were chosen as a bait and the interaction with the juxtamembranous region of TrkB (aa 454-465) as well as with the first 23 aa of TK domain (TrkB-537-559) was verified. With a series of combinations of several regions, the interacting motif of SIPA1L2 was narrowed down. The intensity of the binding in **c,d,f** is represented according to the time it took for the blue staining to be noticeable as following: “+++” <60 min, “++” 60-120 min, “+” 120-180 min, and – for no interaction **g.** GFP-SIPA1L2-δ86-99 shows a significantly decreased interaction with TrkB when compared to fl-SIPA1L2 in heterologous co-immunoprecipitation experiments. **h.** Representative images and line profiles showing the colocalization between fl-TrkB-GFP with SIPA1L2-mCherry, but not with SIPA1L2- Δ14-mCherry. **i,j.** Cartoon representing the time line of the experiment in Fig. **2i,j** and the infusion location (red dot). Images (**j**) show coronal brain sections from injected mice after behavioural tests. Scale bar is 200 µm.

**Figure S5.**
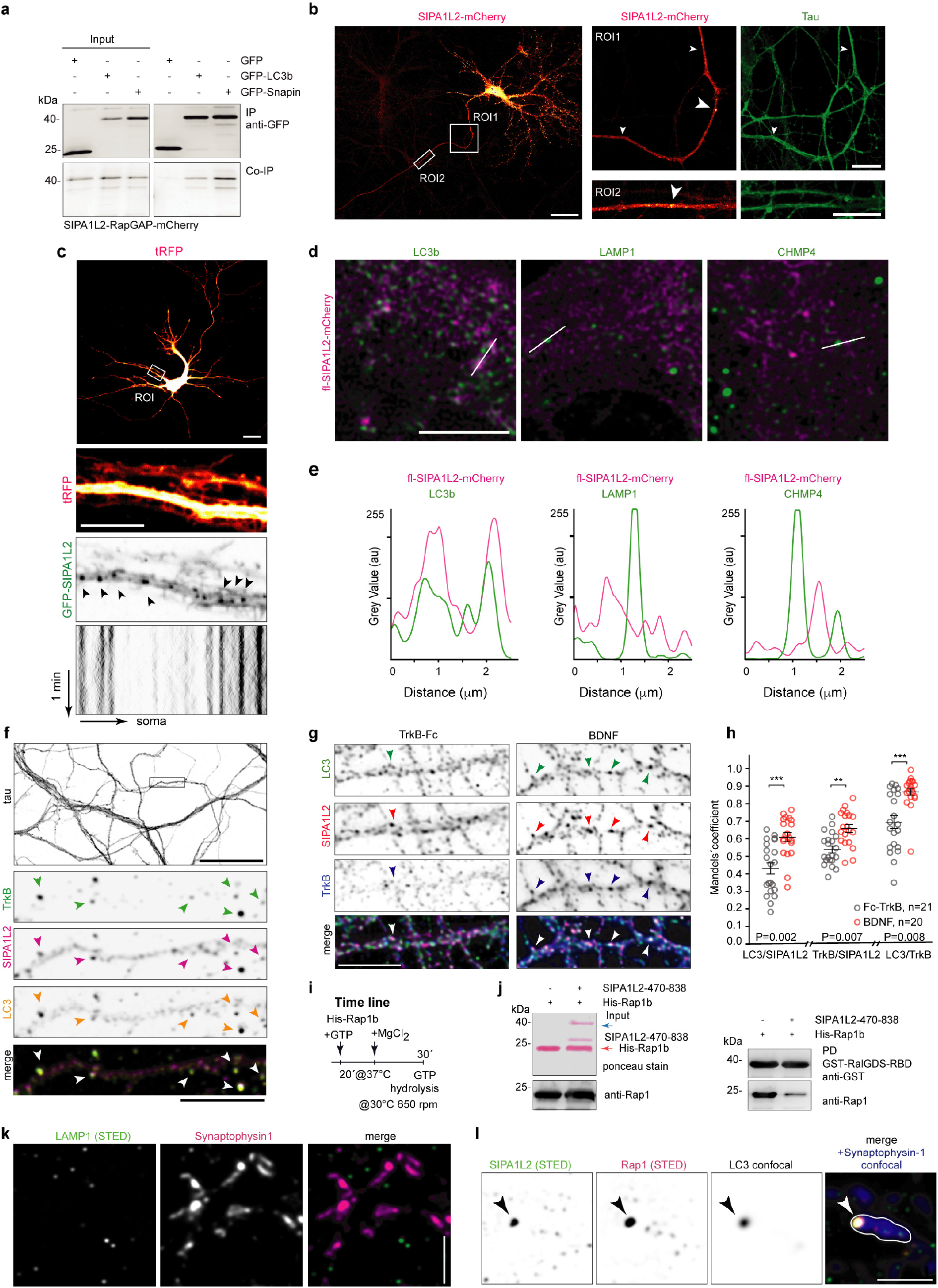
SIPA1L2 associates with Snapin, TrkB and LC3b, but not with LAMP1 and CHMP4 **a.** RapGAP domain of SIPA1L2 (aa 624-813) co-immunoprecipitates with both GFP-Snapin as well as with GFP-LC3 from HET293T cell extracts. **b.** Confocal tile scanned image of neuron overexpressing fl-SIPA1L2-mCherry co-stained with tau showing the distal axonal regions used for co-trafficking assays. Scale bars are 50 µm and 10 µm within the ROIs. **c.** Confocal images showing a neuron (DIV 10) overexpressing tRFP and SIPA1L2-GFP. The depicted ROI shows the dendritic region live-imaged for 5 minutes to assess the dendritic trafficking of SIPA1L2. Kymograph below shows stationary particles. Scale bars are µm and 10 µm within the ROIs. **d,e.** Confocal images of MRC5 overexpressing WT-SIPA1L2-mCherry and stained against endogenous LC3b, LAMP1 or CHMP4 (multivesicular bodies marker) and corresponding line profile (**e**). Scale bar is 5 µm. **f.** Axonal co-clustering of LC3, SIPA1L2 and TrkB. Scale bar = 20 µm in overview image, 5 µm in insert. **g, h.** Confocal images from rat primary cultures immunostained for LC3, SIPA1L2 and TrkB after treatment with TrkB-Fc bodies or BDNF and corresponding Manders’ colocalization coefficient (**h**) Circles represent single analyzed images. Black bars are mean±S.E.M. (Mann-Whitney U test). Scale bar = 10 µm. **i,j.** Sketch (**k**) represents the time line of the Rap1b loading with GTP and conditions for Rap1b-GTP hydrolysis in the RapGAP assay after addition of SIPA1L2. SIPA1L2-(470-838) is generated based on sequence alignment with the catalytically active RapGAP-domain of Rap1GAP (Fig. S1c). **k.** STED imaging revealed no detectable staining for LAMP1 at the presynaptic boutons labeled with Synaptophysin1. Scale bar is 5 µm. **l.** Quadruple immunofluorescence combining confocal and super-resolution STED imaging depicting association of Rap1 with SIPA1L2/LC3 positive amphisomes in the presynaptic boutons. Scale bar is 1 µm.

**Figure S6.**
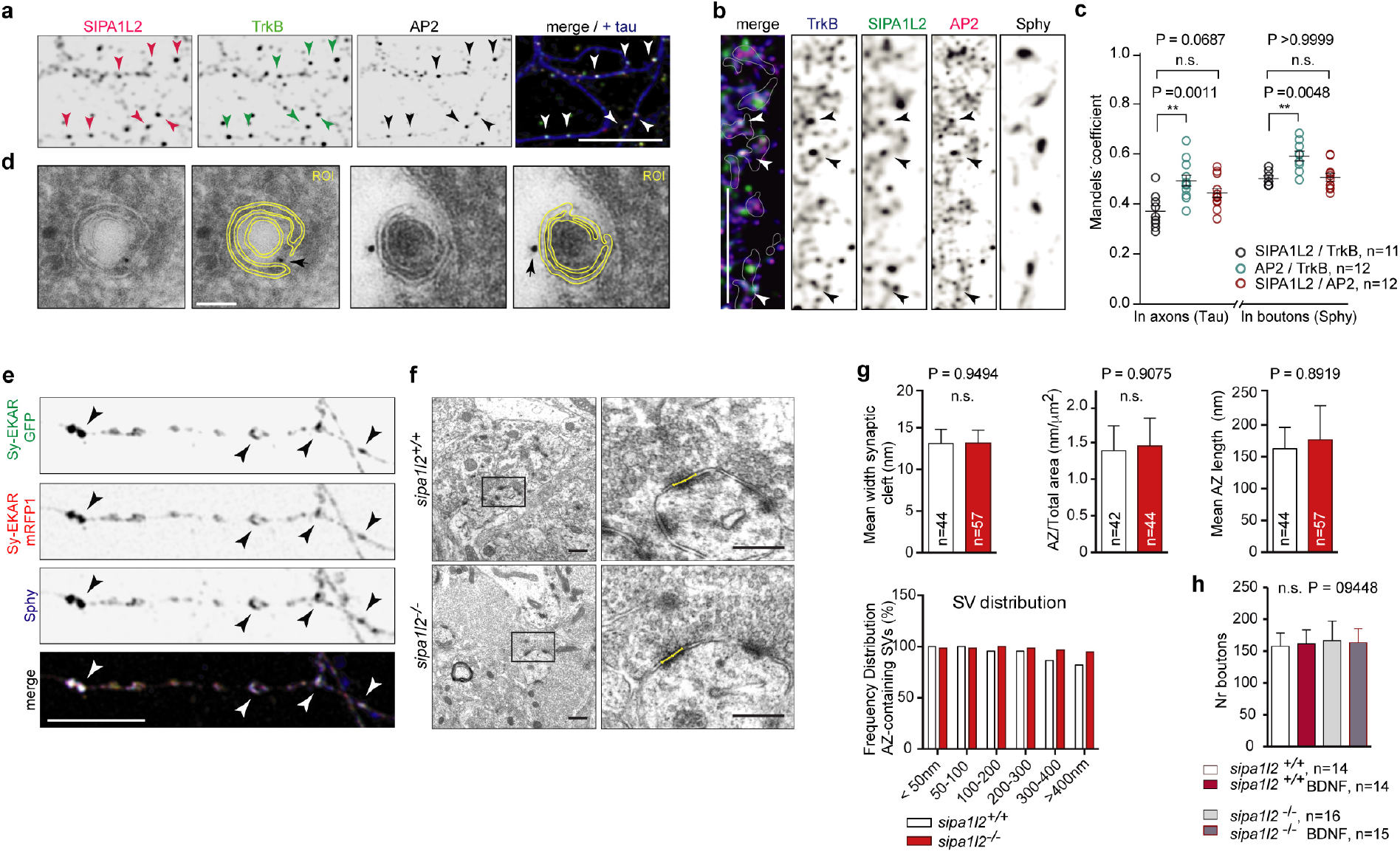
SIPA1L2/TrkB/LC3 signaling amphisomes are positive for AP2 and absence of SIPA1L2 does not result in changes of MF ultrastructure. **a-c.** Confocal images of neurons immunostained with SIPA1L2, AP2, TrkB and tau or Sphy1 and corresponding Manders’ co-localization coefficient (**c**). Scale bars are 5 µm. **d.** Immuno Gold EM image showing the presence of TrkB-tagged at the C-terminus with GFP as part of double membrane organelles in mossy fibre synapses. Note that immunogold particles are located to the outer membrane of these organelles. Scale bar is 100 nm. **e.** Representative confocal images from rat hippocampal neurons transfected with Sy-EKAR (GFP-mRFP1) and immunostained for Sphy-1 showing the localization of the ERK sensor in presynaptic terminals. Scale bar is10 µm. **f-g.** Representative EM images (**h**) and quantification (**i**) of mean width of synaptic cleft, mean active zone length and frequency distribution of AZ-containing synaptic vesicles. Representative active zones are shown in yellow. N numbers are within columns and represent number of images analized. Each image contained at least a MF terminal. Scale bars – 500 nm (overview) and 300 nm (insert). **h.** Number of boutons detected by immunofluorescence against Syn in *wt* and *sipa1l2*^−/−^ neurons after treatment with BDNF as compared to control, non-treated cells. Data shown as mean±S.E.M. “n” numbers represent independent cells. n.s.=p>0.05 (One-way ANOVA with Bonferroni posthoc test).

**Figure S7.**
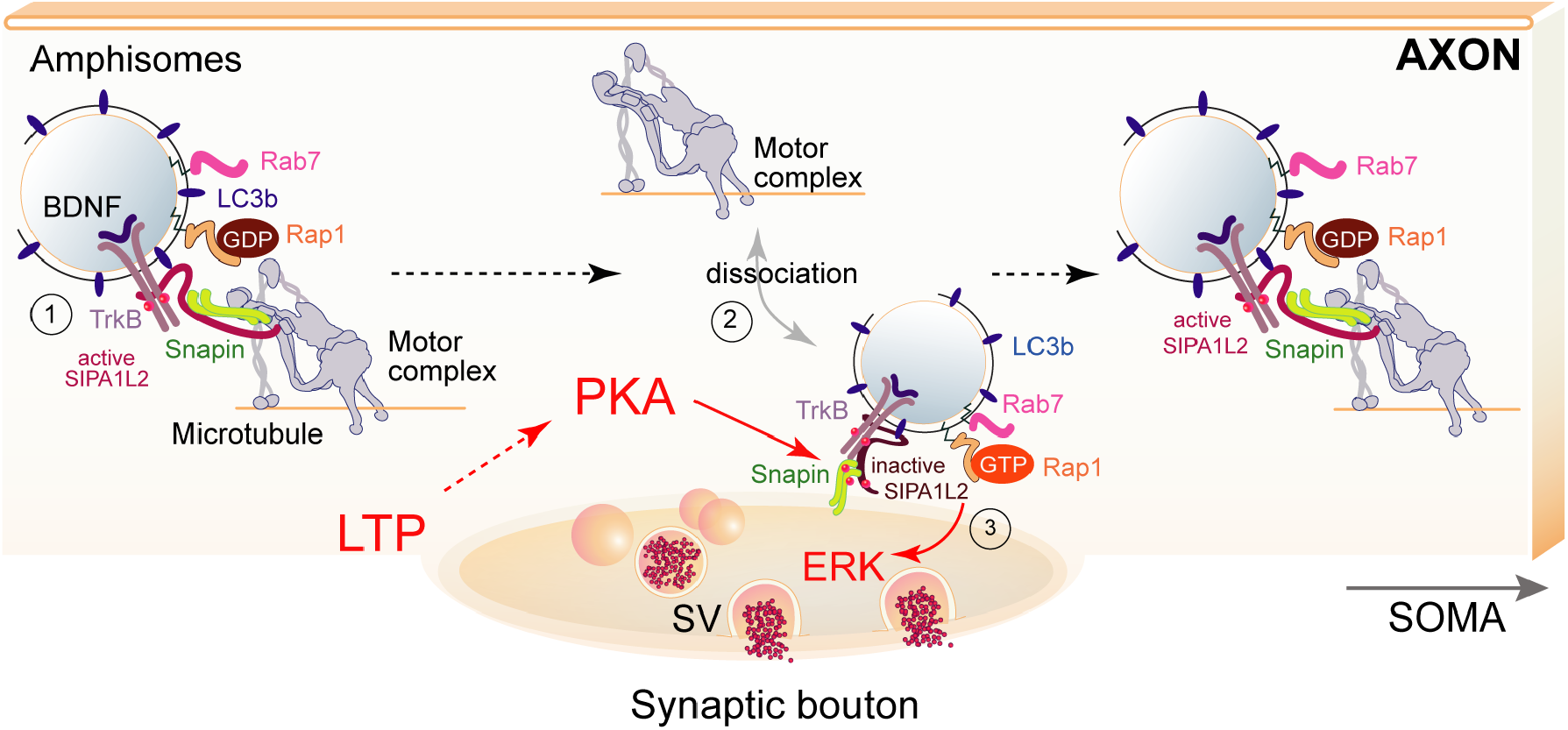
SIPA1L2 assembles a signaling TrkB containing amphisome and controls its retrograde trafficking and signaling at presynaptic boutons (1) Signaling amphisomes result from the incorporation of active TrkB receptors to autophagosomes. Retrograde trafficking and signaling of these amphisomes in axons is tightly regulated by SIPA1L2, which directly binds to TrkB, Snapin and LC3b. While the binding to Snapin serves as a linker to a dynein motor, binding of LC3b enhances SIPA1L2 RapGAP activity which negatively interferes with TrkB/Rap1-signaling and also slows down the velocity of retrograde transport. (2) The amphisome halts at single presynaptic boutons in a PKA-dependent manner by phosphorylation of Snapin, which then dissociates from the motor complex immobilizing the amphisome at individual axonal terminals. (3) PKA activity also terminates RapGAP activity and allows TrkB/Rap1 signaling that subsequently activates synaptic ERK and facilitates transmitter release.

## Materials and Methods

### Animals

Animals were maintained in the animal facility of the Leibniz Institute for Neurobiology, Magdeburg (Germany) or ZMNH, Hamburg (Germany) under controlled environmental conditions. All experiments with animals were carried out in accordance with the European Communities Council Directive (2010/63/EU) and approved by the local authorities of Sachsen-Anhalt/Germany (reference number 42502-2-1264 LIN and 42502-2-1284 UNIMD) or of the city-state of Hamburg (Behörde für Gesundheit und Verbraucherschutz, Fachbereich Veterinärwesen) and the animal care committee of the University Medical Center Hamburg-Eppendorf.

### Generation of *sipa1l2*^−/−^ mice

For constitutive knockout generation, exon 2 of the *SPAR2/sipa1l2* locus (Gene ID: 244668; RefSeqs NM_001081337.1 (mRNA), NP_001074806.1 (protein)) was replaced by a IRESLacZ reporter gene and a neomycin phosphoribosyltransferase selection cassette by homologous recombination. The targeting vector was constructed by amplifying homologous regions from 129S6/Sv/Ev mouse genomic DNA using primers described in Suppl. Table 1 (Fig. S2a,b).

Gene targeting was performed using ES cells (CCB; 129S6/Sv/Ev strain) and chimeras were generated by injection into C57/Bl6 blastocysts. All experiments described here were performed with mice backcrossed for at least 10 generations to C57BL/6J background. Genotyping was performed described in Suppl. Table 1 resulting in bands of 200 bp for wt and 400 bp for ko condition

### Antibodies

A list of antibodies used in this study and corresponding dilutions is available in Suppl.Table 1.

### Cloning

A list with the main contructs used in this study and their source is available at Suppl. Table 1. For Snapin knockdown experiments, a published Snapin KD sequence (64) was cloned into the pSIH-H1 shRNA vector (System biosciences). Inserted sequences are described in Suppl. Table 1.

The Sy-EKAR vector was created by PCR amplification of EKAR-GFP/RFP (38; Addgene #18680) to which a 4Gly linker domain (GGTGGCGGTGGA) was incorporated in the N-terminus (see Suppl. Table 1). This was subcloned into the CMV: ratSyGCaMP2 (Addgene #26124), from where GCaMP2 was removed. SIPA1L2 mutants were generated using the Q5 site directed mutagenesis kit (NEB).

### Yeast Two-Hybrid

The Yeast-Two-Hybrid assay was performed as described in (65).

### Electrophysiology in acute hippocampal slices

#### MF LTP

Experiment was performed as described in (5). Hippocampal slices from 11-16 weeks old C57BL/6J or *sipa1l2*^−/−^ mice were cut and pre-incubated for 3 hours prior recordings. When required, slices were pre-incubated for 3 hours before recording in carbogenated ACSF either with 1 µM of TAT-SIPA1L2 peptide or with 1 µM of TAT-scrambled peptide as a control, as well as with or without 5µg/ml TrkB-Fc.

#### Post-synaptic DG LTP

Field-EPSPs were measured with an electrode positioned at the middle of the molecular layer of the DG. The medial perforant path was stimulated with biphasic constant current pulses (0.1 ms per half-wave duration). Different theta burst stimulation protocols (TBS) or a HFS protocol were used for induction of synaptic plasticity. The TBS protocols consisted of either one episode of brief presynaptic bursts (10 pulses of 0.2ms per half-wave duration delivered @10Hz) repeated 10 times @5Hz, two of these episodes (inter-episode interval 10s) or 4 episodes (inter-episode interval 10s) repeated 4 times every 5min. The HFS protocol consisted of 4 trains of 100 pulses (0.2ms per half-wave duration) @100Hz, inter-train interval 5min.

### Behaviour

#### Spatial Pattern Separation

The behavioral spatial pattern separation task was performed as described (14; Fig. 1c) with minor modifications.

#### Object location and novel object recognition

Two equal objects were placed at equidistant position from the walls of the arena, and the animals were free to explore them. Twenty-four hours later, novel location recognition was assessed as one of the same familiarized objects (A) was displaced to a new location and mice were again left free to explore both objects over a period of 20 minutes. On the third day, a novel object recognition test took place as one of the familiarized objects was now exchanged for a novel object (B). Animals were again left for 20 minutes to explore the objects before being returned to the home-cage.

#### Rotarod

Animals were tested on an accelerating rotarod apparatus 2 trials per day, for 5 consecutive days, in which the speed of the rod was gradually increased each day (from 15 rpm to 36 rpm). On day 5, each animal was submitted to 4 trials lasting 300 sec each, during which the rotation speed gradually increased from 4 to 40 rpm. The latency to fall was recorded.

#### DG infusion of the SIPA1L2 TAT-peptide

Animals (*wt*) underwent sample phase training of the spatial pattern separation protocol, and 5 min after the session received an intra-DG infusion of either 1mM TAT-SIPA1L2 peptide or TAT-scrambled conjugated with fluorescein. Anesthesia was induced at 5% isofluorane in O_2_/N_2_O mixture (Rothacher Medical GmbH., Switzerland) and mice were placed in the stereotaxic frame (World Precision Instruments). During surgery, anesthesia was maintained at 1.5%-2.0%. Craniotomy was done targeting the dorsal dentate gyrus with stereotaxic coordinates (66), anterioposterior (AP) −2.0 mm, mediolateral (ML) ±1.4 mm from Bregma and dorsoventral (DV) −1.6 mm (from brain surface). Each mouse received 1.5 µl bilateral infusion of a respective peptide at a flow rate of 0.5 µl/min. The effect of TAT-peptide infusion was assessed 24 h later in choice phase of pattern separation task. After the completion of the task, mice were deeply anesthetized and perfused transcardially with PBS and 4% paraformaldehyde (PFA) in PBS solution in order to verify DG localization of injections (Fig. S4i,j).

### Cell culture and immunostaining

#### Cell line and primary neuronal culture and transfections

Cell line maintenance and transfection of MRC5, HEK293T and COS-7 cells was performed as described in (67). Primary rat (Sprague Dawley, E18) hippocampal cultures were prepared as previously described (68,69). Mouse hippocampi were dissected and prepared previously described (70) and neurons were plated at a density of 35.000 cells/18 mm-coverslip and maintained as described.

For live imaging experiments primary hippocampal cultures were transfected at 10-13 DIV using Lipofectamin 2000 (Thermo Fisher Scientific) according to manufacturer’s instructions and imaged 48h after transfection. For knock-down experiments, neurons were transfected at 8DIV and imaged 96h after transfection.

When required, BDNF (Tocris, 100 ng/ml) was added into the growing media for 30 minutes before fixing the cells. TrkB-Fc bodies (Acro Biosystems) were added at a concentration of 500ng/mL for 12-16h.

#### Immunocytochemistry

Immunocytochemistry was performed as described in (67,71). Sequential labeling steps were included to avoid cross-labeling. For detection of SIPA1L2, a heat-based antigen retrieval protocol was used. This included the immersion of the cells into a10 µM sodium citrate solution (pH=9) for 30 min at 80°C before proceeding with the standard protocol. The pH of the sodium citrate (Fluka) solution was adjusted with 5M NaOH..

#### Cryosections and staining

Brains of adult *sipa1l2* wt and ko mice were post-fixed in 4% PFA and 0.5M sucrose for 24h and 40 µm thick cryosections were cut. For Nissl staining, sections were acidified with 0.05 M acetate buffer (pH 4) for 5 minutes and incubated with 0.05% cresyl violet acetate solution for 10 minutes. Sections were further incubated with 0.05 M acetate buffer (pH 4) for 3 minutes and dehydrated by ethanol steps. For the volumetric analysis of the DG, serial sections were collected throughout the region. Timm staining was performed as described (72).

#### Immunogold and EM analysis

TrkB-GFP was diluted in EP-buffer (125mM NaCl, 5mM KCl, 1.5mM MgCl2, 10mM glucose, 20mM Hepes, pH 7.4) and the electroporation of mouse organotypic slices was performed as described (73). Fixation was done 24 h after electroporation using 4% PFA 0.1 % GA in PBS for 1 hour. The postembedding immunogold labeling was performed as in (74). Biotinylated anti-GFP was recognized with an antibody anti-Biotin followed by 10nm large protein A gold secondary antibody.

For ultrastructural analysis, ultrathin sections were prepared from six adult male animals as described in (75). Quantification was performed as described in (75,76).

### Microscopy and imaging

#### Imaging cell lines and primary neurons and kymograph preparation

MRC5 cells were co-transfected with WT-SIPA1L2-mCherry together with TrkB-GFP, or GFP-Snapin (Fig.3d). For testing the co-clustering with TrkB, WT-SIPA1L2 as well as SIPA1L2-Δ14-mCherry was co-expressed with TrkB-GFP (Fig.S4h). For recruitment assay COS7 cells were co-transfected with RapGAP-PDZ-SIPA1L2-tRFP and GFP-Snapin (Fig.3b,c). Coverslips were mounted on an imaging ring 24 h after transfections and cells were imaged in DMEM-based growing media (see above).

Rat primary neurons overexpressing WT-SIPA1L2-mCherry together with TrkB-GFP, GFP-LC3b or GFP-Snapin were placed in a Ludin chamber (Life Imaging Services) and imaging was performed at 37°C 5%CO_2_ on a VisiScope TIRF/FRAP imaging system from Visitron Systems based on Nikon Ti-E. The system is equipped with the Perfect Focus System (Nikon), Nikon CFI Apo TIRF 100x, 1.49N.A. oil objective, a back focal TIRF scanner for suppression of interference fringes (iLas2, Roper Scientific), and controlled with VisiView software (Visitron Systems). 488 and 561 nm laser lanes were used to excite the respective fluorophores whose florescence was collected through ET 488/561 Laser Quad Band filters to a The ORCA-Flash 4.0 LT sCMOS camera. Time-lapse imaging of neuronal axons were recorded at 0.5Hz for 4 min.

Analysis was performed using the free software Fiji from ImageJ (http://imagej.nih.gov/ij/) and kymographs were created using KymographClear (77) by drawing a line along axons and fourier-filtered kymographs generated by the plugging were displayed in figures. Retrograde and anterograde trajectories were identified by the respective location of the cell soma or axonal tip during imaging. Stationary trajectories were considered as those moving < 3 µm. Axons from three independent experiments were analyzed.

In experiments where Snapin was knocked-down, imaging was performed 96h after transfection on a Leica TCS SP5 system controlled by Leica LAS AF software using HCX PL APO 63×1.40. Areas of ∼60×15µM (512×128 pixels) were scanned using 488, 561 and 633 nm laser lanes, at 37°C and 5%CO2. SNAP-Cell 647 SiR (0.5µM) was added 1h before starting imaging in order to visualize TrkB-SNAP.

Imaged axons were selected according to the following morphological criteria: i) long thin processes growing far from the somatic region with constant width ii) without recurrent bifurcations and iii) lacking protrusions or spines (Fig.S5b).

#### Live imaging of FM_4-64_ unloading

Primary hippocampal cultures (14-16DIV) overexpressing TrkB-GFP or GFP-LC3b were placed into a field stimulation chamber (Warner Instruments) containing two platinum wire electrodes place 10 mm apart and incubated with 10 µM FM_4-64_ (Thermo Fisher) in extracellular imaging buffer (EIB: 119mM NaCl; 2.5 mM KCl; 2mM CaCl_2_; 2mM MgCl_2_; 30mM Glucose; 25mM HEPES in H_2_0 at pH 7.4) containing 10µM AP-5 and 4µM CNQX (78). Dye loading was achieved by delivering pulses for 30sec@20. Unboud dye was washed-out with EIB containing 1mM ADVASEP-7 (Sigma-Aldrich) for 1 min followed by 2 washes with EIB at 37°C. After 10 min, dye unloading was triggered by a train of 900 pulses @10Hz (79) in the presence of AP-5 (50 µM) and CNQX (10µM) and dye unloading was imaged at a frequency of 1Hz for 110 sec. Ten frames were acquired as a baseline before stimulation. Trafficking of either TrkB-GFP or GFP-LC3b was recorded immediately after the stimulation for additional 10 min @1Hz. After that, a second dye unloading protocol was delivered and imaged as described.

Imaging was performed at 37°C 5%CO_2_ on the VisiScope TIRF/FRAP imaging system from Visitron Systems based on Nikon Ti-E described above. Laser lines of 488 and 633 nm were used to activate the respective fluorophores whose florescence was collected ET 488/640 Laser Quad Band filters to a The ORCA-Flash 4.0 LT sCMOS camera. Pulses were evoked by delivering currents of 60mA for 1ms using an isolator unit controlled by a pulse generator (Master-8). Boutons were considered as “visited” when trajectories overlapped with the signal from the FM dye for > 3 vertical pixels in the kymograph while the term “non-visited” was used for those boutons within the same axon where the overlapping signals were lasting < 3 pixels. Circular ROIs were placed in both types of boutons using Times Series Analyzer V3 from Fiji and mean grey values per ROI per frame were calculated from both streams of images acquired during unloading protocols. Data from each ROI was fitted to a one-phase decay from which tau was calculated using GraphPad Prism.

#### Amphisomal trafficking to presynaptic boutons

Primary hippocampal neurons 14-16DIV overexpressing tRFP-LC3b and either SIPA1L2-GFP or SIPA1L2-N705A-GFP were incubated growing media in the presence of an antibody anti-Synaptotagmin1 coupled to Oyster 660 (1:500) for 1-2h at 37°C 5%CO_2_. Imaging was performed for 5min at 1Hz at 37°C 5% CO_2_ on a Leica TCS SP5 system controlled by Leica LAS AF software using HCX PL APO 63×1.40. Areas of ∼60×15µM (512×128 pixels) were scanned using 488, 561 and 633 nm laser lanes. Fluorescence was collected using 3 HyD detectors.

For experiments cLTP induction, neurons in growing media were treated with Rolipram (0.1 µm) and Forskolin (50 µm) for 10 min and cells were imaged after media replacement with EIB (80). When needed, H89 (10 µm) was added for 30 minutes before proceeding with the cLTP induction.

time of the amphisome in boutons was calculated from kymographs by drawing a vertical line using Fiji in cotrajectories that overlapped with the synaptic marker and number of pixels was quantified. Only those stopovers where the overlap occurs for > 3 vertical pixels were considered. Velocity was calculated from the angle of each cotrajectory obtained by drawing a line along each trajectory. Run length was also assessed by drawing a line and measuring its dimensions using Fiji. Frequency distributions were generated in GraphPad Prism.

#### pSyn/Syn ratio

Hippocampal primary cultures from wt and *sipa1l2*^−/−^ mice were transfected at 10DIV with either WT-SIPA1L2-mCherry, SIPA1L2-Δ14-mCherry or empty-tRFP for control. Transfected neurons were treated at 14DIV with TrkB-Fc (500ng) overnight or with BDNF (100ng) for 30 min.

Immunocytochemistry was performed with anti-Synapsin I,II, anti-phospho Synapsin (Ser62) and anti-mCherry primary antibodies. Stack images were acquired in a Leica TCS SP5 system controlled by Leica LAS AF software using HCX PL APO 63×1.40. Areas of 82×82µM were scanned with 488, 568 and 635 nm laser lanes (12-bits, 80×80nm pixel size, 700Hz, Z-step 0.25µm). Fluorescence was collected using 3 HyD detectors.

Circular regions of interest (ROIs) were placed using Time Series Analyzer v3 on the Synapsin signal and pSyn / Syn ratio was calculated for single boutons from the mean grey intensity obtained in Fiji for each ROI. For each rescue group, data was normalized to the group of neurons treated with TrkB-Fc.

#### Imaging of Sy-EKAR

Neurons overexpressing Sy-EKAR and either with SNAP-SIPA1L2 or SNAP-LC3b were imaged in EIB. The far-red SNAP-Cell 647 SiR substrate at a concentration of 0.5 µM was added to the growing media for 1h and was washed out (3X) with EIB.

FLIM measurements were carried out at 37°C and 5%CO_2_ using an Abberior multi-channel confocal/STED microscope based on a Nikon Ti-E microscope body with perfect focus system. For excitation 3 pulsed lasers at i.e. 470, 561, 640 nm were employed. A 60x P-Apo oil immersion objective (NA 1.4) was used. In the detection path the emitted light was directed through a pinhole set to match 1 Airy at 540 nm. Three avalanche photo diodes APDs were used for recording the fluorescence signal in the green 470-520 nm red 595-635 nm and far red 615-755 nm range. For measuring the fluorescence lifetime a Becker&Hickl SPC150 TCSPC board was used after setting the APDs in single photon count mode (spcm).

A small area of interest containing 2-5 boutons was chosen. Two hundred FLIM images were recorded in streaming mode detecting only the green and the far red channel. In this way it was possible to record 2-4 FLIM images/sec (pixel size was set to be 110 nm). The acquisition speed of any FLIM time-lapse experiment carried out in this work can be calculated using this formula:

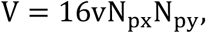

where V is the acquisition speed in frames/s, 16 is the total number of line accumulations used for imaging the green and far red channel, V is the pixel dwell time set to be 3 µs in all the experiments and N_px/y_ is the x and y axis image pixel size. FLIM measurements were carried out sampling the acquisition period of 19 ns in 50 steps of 0.38 ns.

FLIM images were analysed by a self-made Matlab script where boutons were automatically localized by finding the local peaks of the intensity signal in the sum projection images and 5×5 pixels-ROIs were place. The time series of the mean fluorescence lifetime was produced from each ROI by averaging the pixel by pixel measured lifetime. The life time information was extracted with the help of a least square algorithm. A mono exponential decay function was used as model function in the least square algorithm.

Boutons in axons without any detected trafficking were used as a control. Mean lifetime calculated from each bouton before - 4 frames-, during and after the visit of the amphisome and ΔLifetime was calculated for each bouton. Lifetime_0_ was considered as the mean lifetime in the frame before the arrival of the amphisome.

#### Confocal and STED Imaging

Gated STED images were acquired with a Leica TCS SP8 STED 3X equipped with pulsed White Light Laser (WLL) and diode 405 nm laser for excitation and pulsed depletion with a 775 nm laser. A Leica HC APO CS2 100x/1.40 oil objective was used. Images were taken as a single plane of 1024×1024 pixels and optical zoom ranging from 5 to 6, at 400 lines per seconds and 4x line averaging. Pixel size was 20-23nm in xy. Time gates were set to 0.5 to 6 ns. Raw STED and confocal images were deconvoluted using Deconvolution wizard (Huygens Professional, SVI) where using a theoretical point spread function (PSF) based on optical microscopic parameters until it reached a Quality threshold of 0.05.

Mander’s coefficient was calculated on deconvoluted images using either co-localization analyser plug-in (Imaris, Bitplane) or the Colocalization test plugin from Image J. Briefly, region of interests (ROIs) for presynaptic boutons were defined by intensity threshold of Bassoon staining. Manders M1 coefficient (SIPA1L2/ pTrkB^Y515^) with Costes thresholding applied (81) was measured within the synaptic boutons as a fraction of the total fluorescence found in common presynaptic ROIs.

### Biochemistry

#### Co-immunoprecipitation

Co-immunoprecipitation experiments were performed as described before (65). Briefly, cerebellar tissue was homogenized in RIPA buffer (20 mM Tris, pH 7.5, 150 mM NaCl, 1% Triton X-100, 1% sodium deoxycholate, 1 mM EGTA and 1x protease inhibitors), Guinea pig anti-SIPA1L2 sera, 5 µg of anti-TrkB or 5 µg unspecific IgG were incubated and covalently cross-linked with Protein-A sepharose beads (Amersham) that were washed with RIPA buffer and collected by centrifucation. The IgG-coupled beads were added to cerebellar extracts and incubated overnight @4 °C with rotation. The unbound fraction was removed and sepharose was washed in extraction buffer. Proteins were eluted and subjected to SDS-PAGE.

For immunoprecipitation with anti-DIC antibodies, crude light membrane fraction obtained from whole mouse brains was resuspended in homogenization buffer (10 mM HEPES, 320 mM Sucrose and 1 mM EDTA, pH 7.4; PhosSTOP phosphatase inhibitors and complete proteases inhibitors (Roche Holding AG, Basel, Switzerland) containing 1% Triton X-100. 5 µg of anti-DIC antibodies or IgG mouse control antibodies were coupled to magnetic Dynabeads Protein G for 16 h. Subsequently, the coupled beads were washed three times in PBS pH 7.4 containing 0.1 % (w/v) BSA and re-suspended in 200 µl binding buffer (PBS, pH 7.4, containing 2 mM EDTA and 0.1% (w/v) BSA. For co-immunoprecipitation, 120 µl of the crude light membrane fraction was added and incubated for 16 h with constant rotating. All steps were performed @4°C. After substantial washing, proteins were eluted from the beads in 60 µl SDS sample buffer, boiled at 95°C, subjected to 8 % SDS-PAGE and analyzed by WB.

#### Lipid raft isolation

Cerebella were homogenized in lysis buffer (50 mM Hepes pH 7.4, 100 mM NaCl, 3% Brij 58, 5 mM EDTA, 10 mM Na_4_P_2_O_7_, protease inhibitors). The lysate was mixed with 1.5 ml ice-cold sucrose (final concentration: 40%) and transferred to a Potter S (B.Braun, Germany). A homogenization step was performed and 3 ml of homogenate were transferred into a centrifuge tube. The sample was overlayed with 6 ml ice-cold 30% sucrose and then again overlayed with 3 ml ice-cold 5% sucrose and subjected to centrifugation for 20 hours at 200,000 x g @4°C. Twelve samples of 1 ml obtained and proteins of each sample were precipitated with acetone. Fraction 13 (pellet) was diluted in lysis buffer. Pellets from all fractions were lyophilized and resuspended in SDS protein sample buffer for SDS-PAGE and immunoblotting.

#### Heterologous co-immunoprecipitation

HEK-293T cells growing in T75 Flasks (Thermo Fisher Scientific) were transfected, harvested and lysed in 1ml of RIPA buffer (50 mM Tris-HCl pH 7.4, 150 mM NaCl and 1 % Triton X-100, 0.5% sodium deoxycholate and 0.1% sodium dodecyl sulfate (SDS)) for 1h@4 °C. The lysate was cleared by centrifugation incubated with anti-GFP/myc antibody coated magnetic beads (MultiMACS GFP Isolation Kit #130-094-253 or Miltenyi Biotec GmbH, Germany). The co-immunoprecipitation was done following the manufacturer’s protocol (Mitenyibiotec, Bergisch-Gladbach, Germany). For co-immunoprecipitation of WT-SIPA1L2-mCherry and SIPA1L2-δ14-mCherry with TrkB-GFP the following lysis buffer was used: 50 mM TRIS, 150 NaCl, 1% NP-40, 0,1% SDS, protease inhibitors (Complete, Roche), PhosphoStop (Roche). For heterologous co-immunoprecipitation of RapGAP and PDZ domains of SIPA1L2 with GFP-Snapin as well as co-immunoprecipitation of endogenous DIC and SIPA1L2-mCherry with either GFP-Snapin-S50A or GFP-Snapin-S50D the lysis buffer was without SDS (Fig.3a).

#### Pull-down assays

Proteins were bacterially purified as previously described (65).

For interaction assays between SIPA1L2 and TrkB, the matrix (amylose-MBP) alone or coupled with different fragments of the TrkB receptor was washed three times and resuspended in pull down buffer (20 mM Tris pH 7.5, 1 mM DTT, 3 mM EDTA, 100 mM NaCl, 0.3% TritonX-100) and protease inhibitors (Complete, Roche). Subsequently, amylose resin was incubated with 2µg of a GST-ActI-SIPA1L2-1-624 or GST-ActII-SIPA1L2-1026-1650 overnight @4°C. Reciprocally, 10 µg of ether GST-SIPA1l2-86-99 or GST alone as a control was immobilized on Glutathione Sepharose resin equilibrated with PBS buffer (140 mM NaCl, 2.7 mM KCl, 10 mM Na_2_HPO_4_, 1.8 mM KH_2_PO_4_, pH 7.3, protease inhibitor (Complete, Roche), incubated with 200 µg of purified MBP-TrkB-454-465 for 120min@4°C in PBS buffer and washed 3x times with PBST buffer (PBS, protease inhibitor, 0.1% Triton X-100). All samples were eluted with 2X SDS loading buffer and equal amount of samples were used for SDS-PAGE (Fig.2g., Fig. S4e)

For experiments with the TAT peptides, MBP-TrkB-454-821 was coupled to amylose resin (NE BioLabs) and incubated in the presence of 10x molar access of either TAT-SIPA1L2 or TAT-Scr peptides with 1 µg of GST-SIPA1L2-1-624 fusion protein @4°C overnight. The resin was washed three times with 1 ml of pull down buffer (see buffer composition above) and boiled with 50 µl of 2xSDS sample buffer for 5 minutes prior to SDS-PAGE. Blots were then analyzed by immunodetection with anti-SIPA1L2 antibodies (Fig.2h).

For interaction between SIPA1L2 and LC3, 5 µg of Intein-RapGAP-SIPA1L2 (470-838) and Intein-Caldendrin were immobilized on chitin resin (NE BioLabs). Resin was washed with TBS buffer (150 mM NaCl, 20 mM Tris-Cl, pH 7.4, 5 mM MgCl2, EDTA free protease inhibitors (Complete, Roche) and incubated with purified 6xHis-tag LC3 for 120 min @4°C in TBS buffer either alone or in the presence of purified 6xHis-Snapin. Samples were then washed with TBS-T buffer (TBS + 0.1% Triton X-100), eluted with 2X SDS sample buffer, subjected to SDS-PAGE and immunoblotting (Fig.4d,e).

For Pull Down assay of SIPA1L2 with endogenous LC3 whole brain extract was incubated with GST-SIPA1L2-RapGAP, GST-SIPA1L2-PDZ and well as with GST alone. Immunodetection was done with anti-LC3b antibodies (Fig. 4c).

#### Rap1-GAP activity assay

RapGAP activities were assessed using Active Rap1 detection kit (Cell Signaling) (82) following the manufacturer instructions. For experiments involving overexpression of WT-SIPA1L2, RapGAP and LIR SIPA1L2 mutants GST-RalGDS PD of GTP-bound Rap1 was performed from HEK293T cell extracts (Fig.3g-j). For experiments involving LC3b and PKA phosphorylation of SIPA1L2-S990, Intein-tagged SIPA1L2-470-838, SIPA1L2-470-1025, SIPA1L2-470-1025-S990D, 6xHis-tagged Rap1b as well as 6xHis-targged LC3b were bacterially purified. Purified Rap1b was incubated with 50 mM GTP in 200 µl of loading buffer (50 mM TRIS, 100 mM NaCl, 2mM EDTA, 1 mM DTT) for 20 minutes at 37°C for loading recombinant Rap1b with GTP. The loading reaction was terminated by adding 10 mM MgCl_2_. Subsequently, RapGAP assay was carried out in presence of SIPA1L2-RapGAP domain (470-838aa, ∼20 µg), RapGAP-PDZ (470-1025, ∼30 µg) and RapGAP-PDZ (470-1025-S990D, ∼30 µg) of SIPA1L2 in the presence of either His-tagged LC3b (∼50 µg) or His-SUMO (∼50 µg) as a control for 15 minutes at 30°C on shaker (650rpm) (Fig.3k-m, Fig.S5l,m., Fig.6g-i).

#### Preparation of an autophagosome-enriched fraction

A whole brain homogenate from two rats was subjected to density gradient separation and purification as already described (83). Briefly, rat brains were homogenized and subject to differential centrifugation through Nycodenz gradient (15, 20, 24, 26, 85%) for 3h@105g. Each fraction (A1 −15-20%, A2 - 20-24% and L - 24-26% interfaces) was collected, diluted with Percoll and centrifuged for 1h@105g to obtain a virtually pure pellet and analyzed using antibodies against SIPA1L2, TrkB, Snapin, LC3b and Rab7.

### Statistical analysis

Data in the manuscript is shown as mean ±S.E.M. Graphs and statistical analysis was made with GraphPad Prism (GraphPad Software). Statistical tests used are written in the figure legend of the corresponding experiment. Normality tests were performed to decide between parametric and non-parametric tests. Number of subjects considered for statistical comparison is described in graphs and/or figure legends associated to each experiment. P values <0.05 were considered significant.

---

**Supplementary Table 1.**
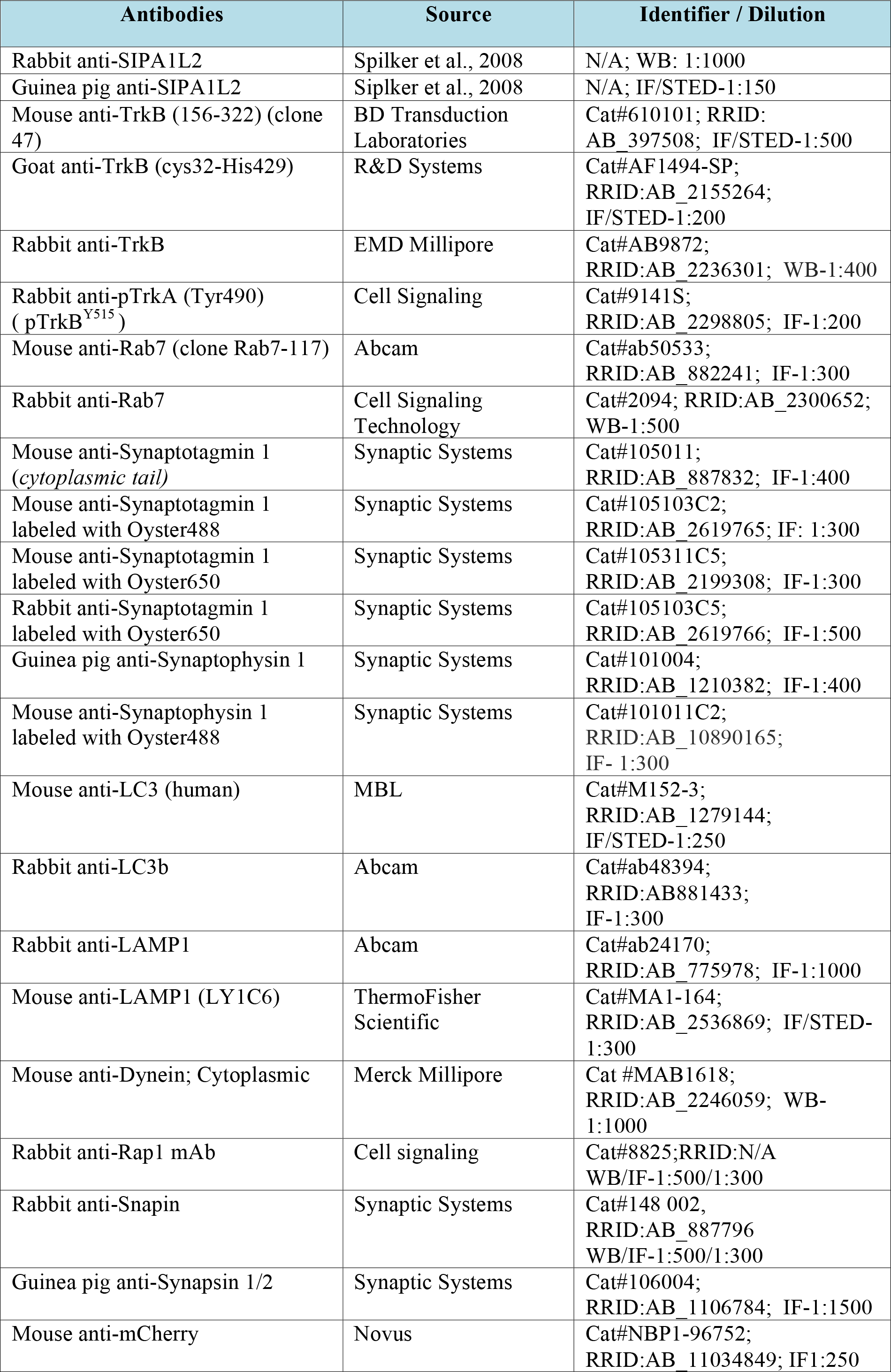
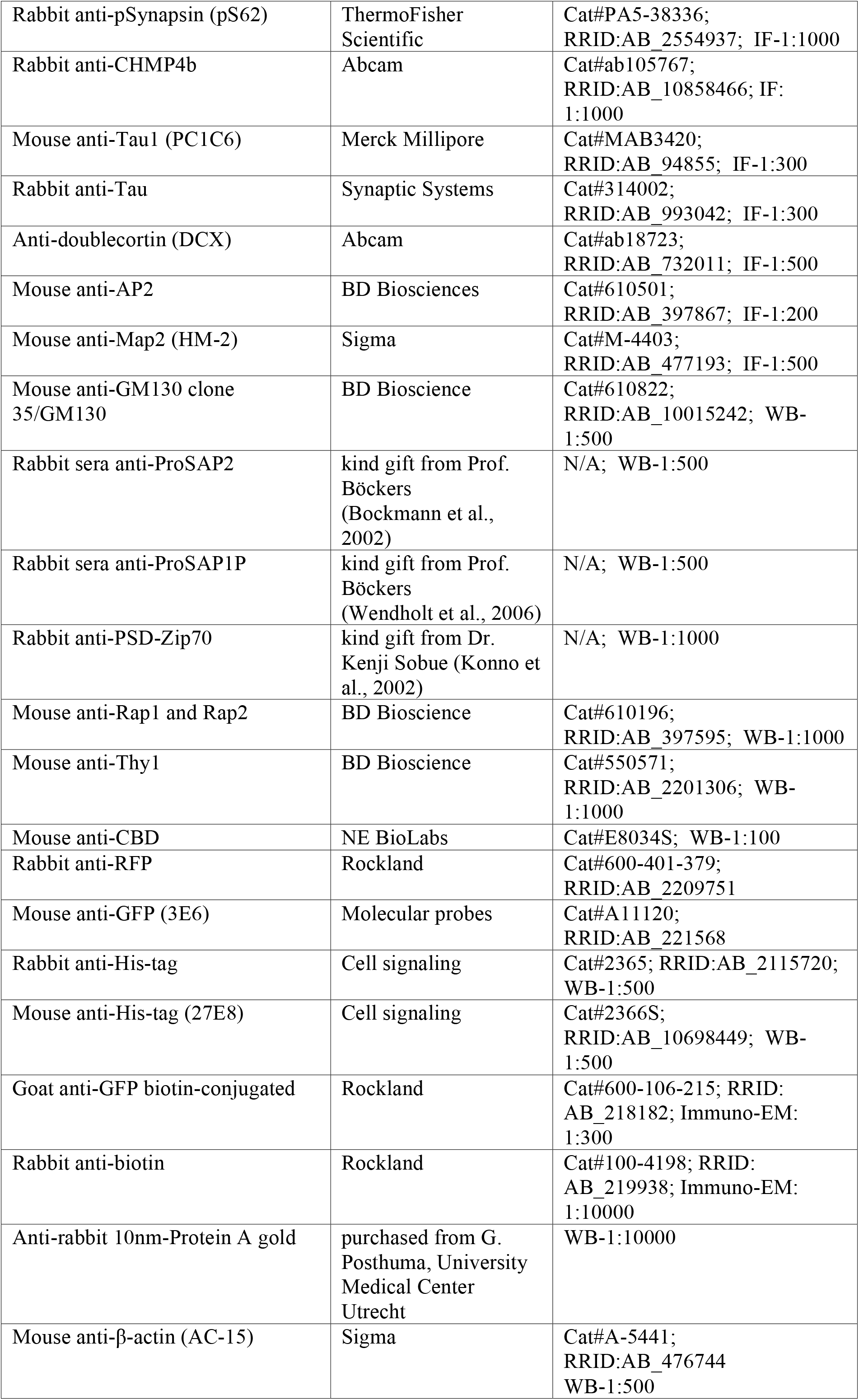
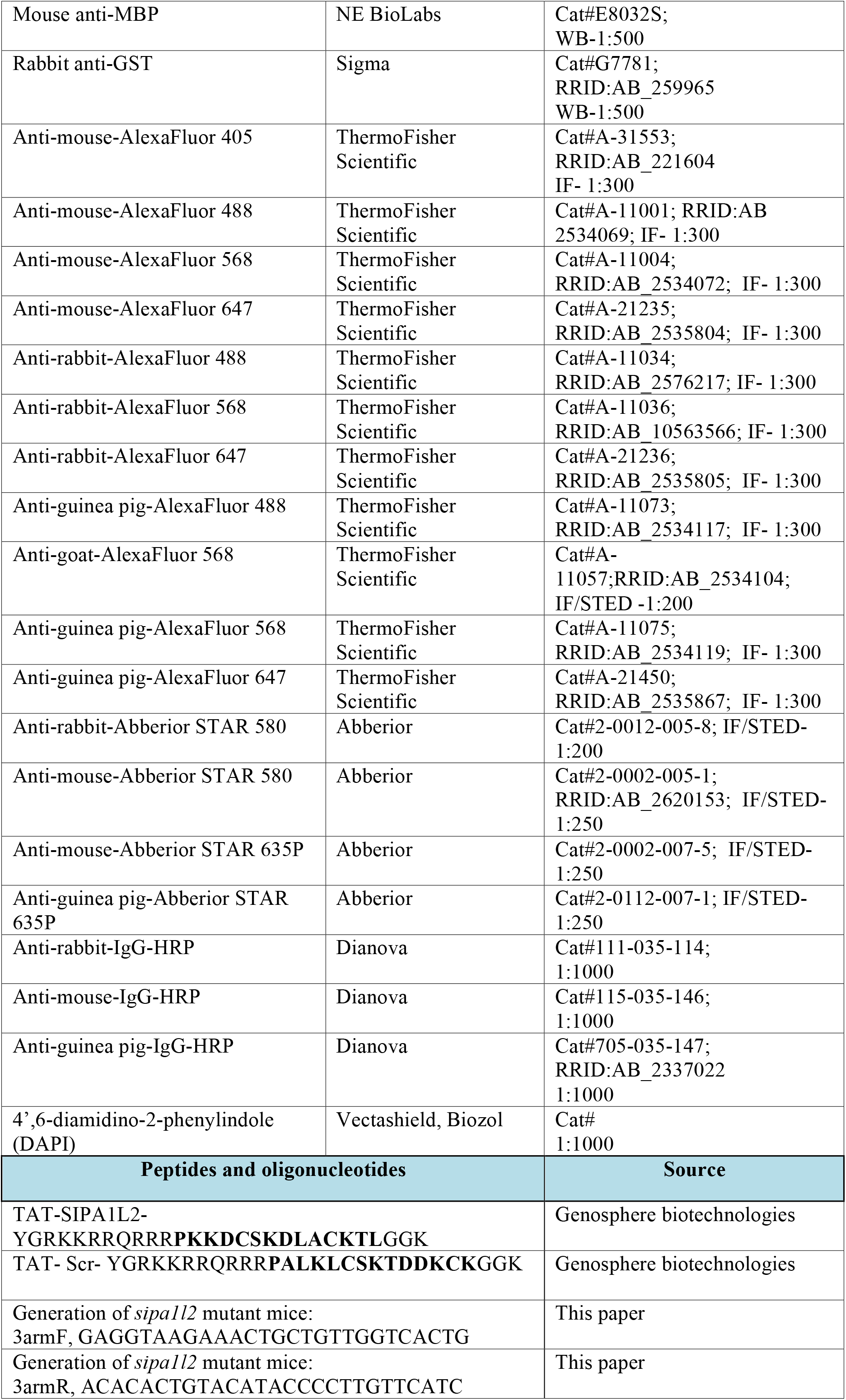
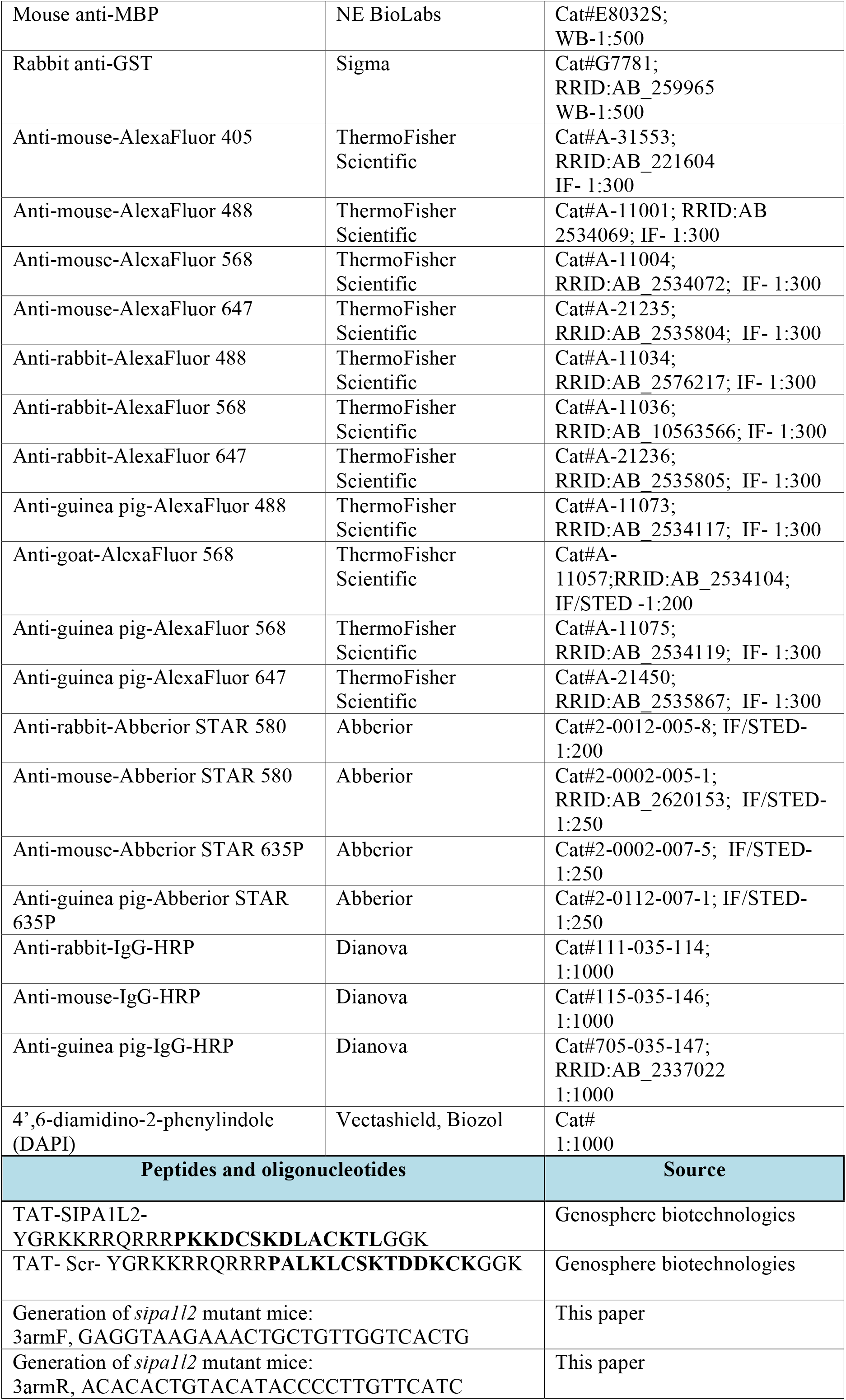
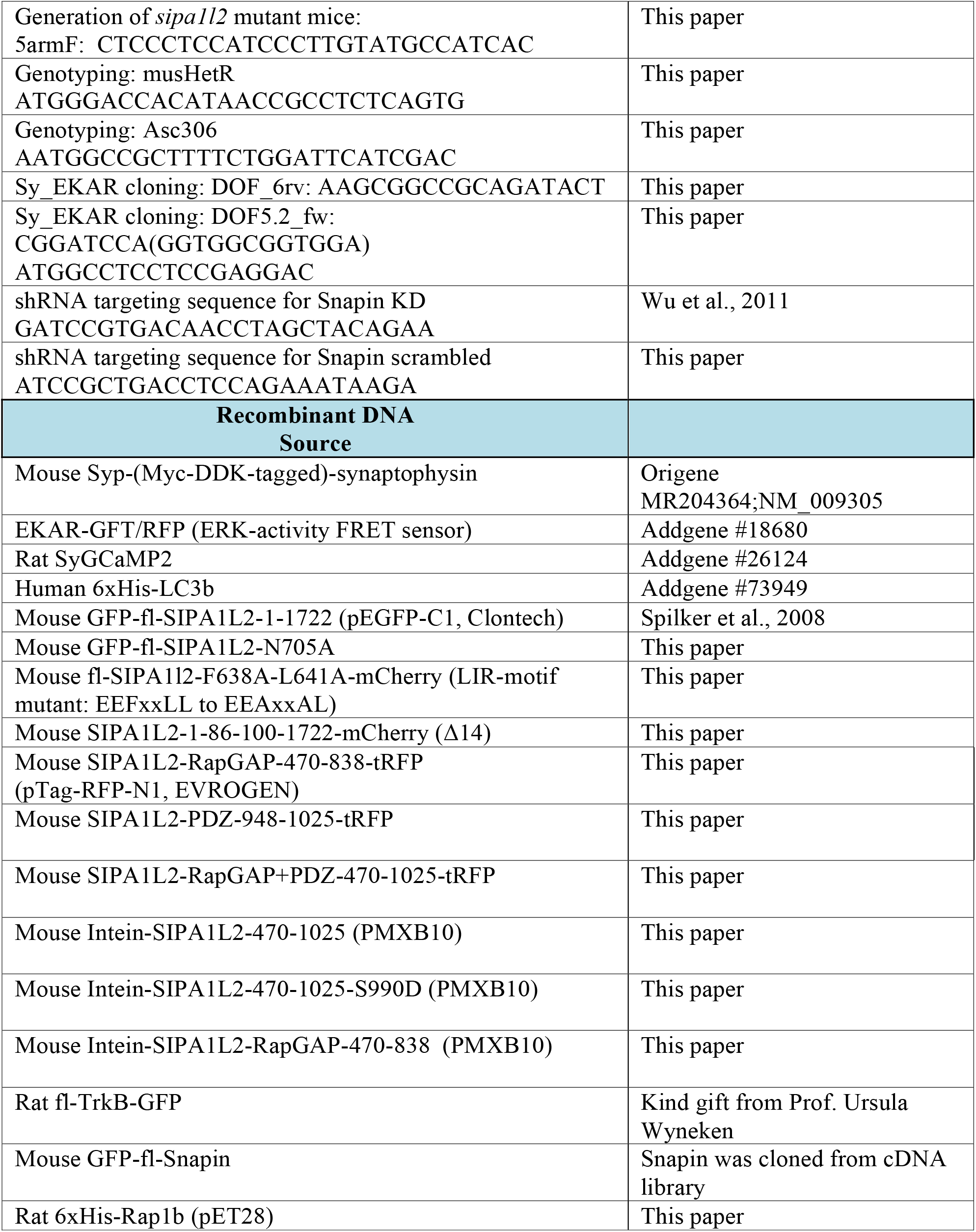
Description of the antibodies, constructs and oligonucleotides used in
this study and corresponding sources.

